# Life-history traits may buffer genetic erosion under isolation in mountain sky-island systems

**DOI:** 10.64898/2026.05.27.727838

**Authors:** Noèmie Collette, Sébastien Pinel, Pascaline Salvado, Maria Martin, Josep Parera, Xavier Oliver, Clara Pladevall, Gerard Giménez Pérez, Ingrid Forey, Anais Gibert, Valérie Delorme-Hinoux, Joris A.M. Bertrand

## Abstract

Sky-island systems provide natural case studies for understanding how geography and Quaternary history shape genomes, phenotypes and life-history traits in mountain endemics. We investigated *Xatartia scabra* (Apiaceae), a monotypic scree specialist plant species restricted to sky-island summits in the eastern Pyrenees, by integrating population genomics with abiotic condition-informed distribution modeling. Using a ddRAD-seq-like protocol (nGBS), we genotyped 125 individuals (21,970 SNPs), and applied species distribution modeling to identify suitable environmental conditions from the Last Glacial Maximum to 2100.

Genetic analyses revealed unexpected genetic “resilience”, with moderate genome-wide diversity (*H*_E_ = 0.17, Ho = 0.15) and low inbreeding (*F*_IS_ = 0.07), despite small census sizes and strong isolation. Significant overall genetic differentiation (*F*_ST_ = 0.16), with pronounced summit-level structure and strong isolation-by-distance, supports deep valleys as barriers to gene flow.

Abiotic niche reconstructions recovered extensive Heinrich Stadial 1 connectivity (+ 58,4% relative to present), followed by postglacial loss of suitable habitats at lower elevations and increasing fragmentation; projections under climate change forecast contraction (−61,2%, relative to present) of climatic suitability, expected to intensify drift in small, isolated populations. Demographic inferences align with this narrative, indicating postglacial decline in effective population sizes.

Taken together, genetic, climatic, and demographic evidence supports a transition from historically connected lowland corridors to modern sky islands where distance-limited drift dominates. The maintenance of moderate genetic diversity and low inbreeding under strong isolation suggests that *X. scabra* may have evolved life-history strategies including outcrossing and monocarpy that allow limiting genetic erosion in such extreme and fragmented environments.

## Introduction

Mountains represent reservoirs of biodiversity and endemism worldwide (Rahbek *et al*., 2019b). Their steep environmental gradients, topographic complexity (Marder *et al*., 2025), and climatic heterogeneity (Rahbek *et al*., 2019a ; 2019b), together with Quaternary glacial-interglacial cycles, have historically promoted diversification and long-term species persistence (Harrison & Noss, 2017 ; Nevado *et al*., 2018). In contrast, contemporary warming is occurring at an unprecedented rate and magnitude, leaving little time for species to track changing conditions, turning the features that once fostered diversification into drivers of vulnerability (Chan *et al*., 2024 ; Rumpf *et al*., 2019 ; Trew & Maclean, 2021). This vulnerability is particularly pronounced in mountain sky-island systems.

By analogy with true islands (water-surrounded islands), spatial isolation and habitat limitation are expected to be associated with small population sizes and restricted gene flow in such systems, resulting in genetic erosion through reduced diversity (Pinto *et al*., 2024 ; Rubidge *et al*., 2012). Such conditions promote genetic drift and inbreeding (Ellstrand & Elam, 1993 ; Maruyama & Fuerst, 1985), which in turn reduce adaptive potential and increase extinction risk, particularly under rapid environmental change (Bijlsma & Loeschcke, 2012 ; Frankham, 2005 ; Mathur *et al*., 2023). Consistent with this analogy, mountain sky-island systems are therefore expected to experience strong demographic and genetic consequences under ongoing climate change (Love *et al*., 2023). While these patterns are well documented in island systems (Franks, 2010 ; Hamabata *et al*., 2019) and increasingly reported in mountain contexts (Hartley *et al*., 2023 ; Zhang *et al*., 2019), fewer studies explicitly link projected habitat changes to contemporary genetic structure within a predictive framework.

These expectations are particularly relevant for mountain endemics, which represent extreme cases of spatial isolation within alpine systems. Shaped by long-term geographic isolation, refugial persistence, ecological specialization (Harrison & Noss, 2017 ; Love *et al*., 2023 ; Nery *et al*., 2023 ; Veron *et al*., 2019), these species frequently occur in small, isolated populations. While contributing to regional biotic uniqueness (Lamoreux *et al*., 2006), such conditions increase vulnerability to environmental change (Manes *et al*., 2021) and demographic stochasticity (Ellstrand & Elam, 1993). Quantifying the genetic consequences of isolation in these systems is therefore essential for predicting their responses to ongoing climate change.

The case study presented here combines all three dimensions: endemism, isolation, and small population size. *Xatartia scabra* (syn. *Xatardia scabra*) (Lapeyr.) Meisn. (Apiaceae) is the sole representative of its monotypic genus (Clarkson *et al*., 2021). This micro-endemic plant is restricted to the eastern Pyrenees, where it occurs on ten isolated summits within a total longitudinal extent of less than 120 km. Its habitat consists of unstable limestone and schist screes (Huc, 2008) between 1,600 and 2,800 m above sea level. Biologically, *X. scabra* is a perennial hemicryptophyte, monocarpic and hermaphroditic. It is entomophilous and strictly allogamous due to self-incompatibility, and reproduction is followed by seed dispersal by both barochory and anemochory (Huc, 2010). Cytogenetic and flow cytometry analyses report a chromosome number of 2n = 22 and 2C-value = 9.1 pg, corresponding to a haploid genome size of 4.5 Gbp for this species (Centre Nacional d’AnÀlisi GenÒmica, 2025). *X. scabra* thus embodies a unique evolutionary lineage whose loss would not only diminish local biodiversity but also see disappear witness to the historical biogeography of the Pyrenean region and sky island systems in general (Vargas, 2023). Preserving phylogenetically distinct taxa is recognized as a goal in conservation prioritization, as it safeguards evolutionary potential (Cardillo, 2023 ; VÉzquez & Gittleman, 1998 ; Winter *et al*., 2013), a component to cope with accelerating global change. In this context, *X. scabra* represents both a pressing regional conservation challenge and a valuable model for understanding the persistence of evolutionary distinct species in rapidly changing environments.

Despite its ecological and evolutionary singularity and its protected status across France, Andorra, and Spain (GarcÍa GirÓn & MartÍnez GarcÍa, 2018 ; Hinoux, 2025), little is known about the genetic diversity and the connectivity of current populations as well as the biogeography history of *X. scabra*. In this study, we explicitly test the sky-island predictions in this micro-endemic mountain plant by combining species distribution models (SDMs) and population genomic data. Specifically, we test the expectation that long-term spatial isolation and climatic fragmentation have resulted in (i) low levels of within-population genetic diversity, (ii) significant departures from panmixia and strong population structure, and (iii) reduced effective population sizes. We further evaluate whether past climatic oscillations have left detectable imprints on genetic structure and demographic history, and assess the potential persistence of populations under future climate change. By coupling population genomics data with climate dynamics, our study tries to resolve the interplay of glacial legacies, topographic barriers, and drift-migration balance in mountain endemic species and provides actionable guidance for monitoring and management under rapid warming.

## Materials and Methods

### Population sampling and study area

We sampled 125 *X. scabra* individuals from 11 localities that are representative to the known geographic distribution of this species endemic to the Pyrenees: 1) Mantet; 2) Eyne; 3) Puigmal; 4) Cabaneta; 5) Rialb; 6) Tartera de Ferreroles; 7) Queralbs sector 1; 8) Queralbs sector 2; 9) Castellet de Moro; 10) Tosa d’Alp and 11) Cadí. Twelve individuals were sampled per locality (except for Tartera de Ferreroles, n = 5) (Figure 1, Appendix A). Healthy young leaf tissues (approximately 1 cm²) were excised using sterilized scissors and stored in 1.5 mL tubes filled with 70% ethanol. Back in the lab, the samples were stored at 4° C in 90 % ethanol until DNA extraction to minimize degradation and contamination.

**Figure 1.**
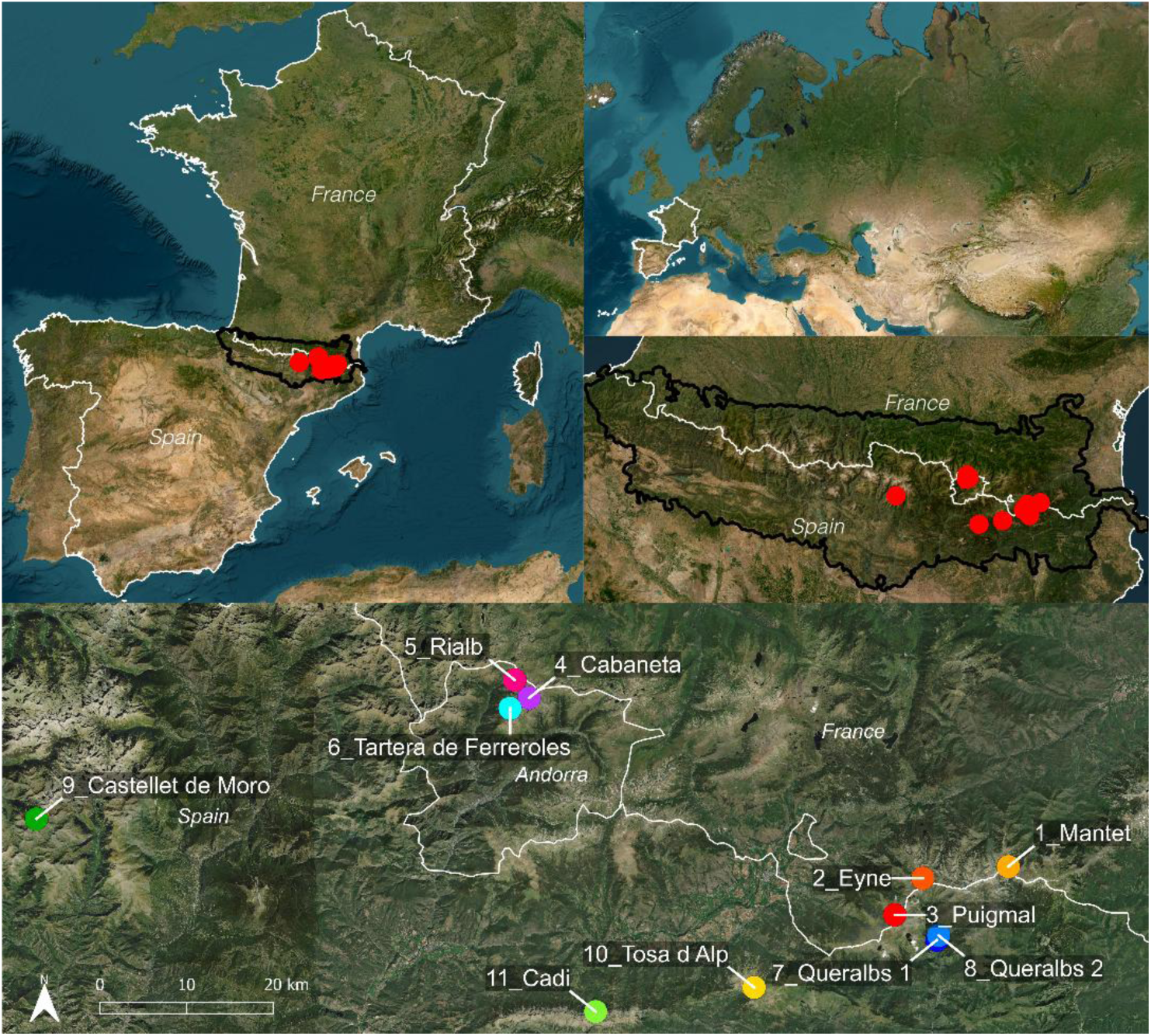
Geographic distribution and sampling design of *Xatartia scabra* populations in the eastern Pyrenees. Geographic distribution of the 11 sampled populations of *X. scabra* in the eastern Pyrenees (France, Spain, and Andorra). The upper panels locate the study area within western Eurasia and within the Pyrenean mountain range. The lower panel presents a detailed map of the eastern Pyrenees showing the sampling locations. Colored points correspond to the sampled populations, and white lines indicate national borders.

### Molecular procedures

Genomic DNA extraction and genotyping were subcontracted to LGC Genomics GmbH (Berlin, Germany). Libraries were generated following an nGBS (normalized Genotyping-By-Sequencing) protocol using a double digest RADseq (ddRAD) strategy. PstI and ApeKI restriction enzymes were employed to reduce genome complexity by targeting specific DNA sites. A normalization step using the restriction enzyme MstI was incorporated to limit the over-representation of highly-methylated repetitive regions. A total of 125 barcoded libraries from *X. scabra* were sequenced in paired-end mode (2 × 150 bp) on an Illumina NovaSeq 6000 platform. The data for this study have been deposited in the European Nucleotide Archive (ENA) at EMBL-EBI under accession number PRJEB113243 (https://www.ebi.ac.uk/ena/browser/view/PRJEB113243).

Reads were processed using Stacks v.2.62 (Catchen *et al*., 2013). After demultiplexing and quality filtering, loci were assembled *de novo* (*ustacks*) and combined into a catalog (*cstacks*) against which all individuals were matched (*sstacks*). Single Nucleotide Polymorphisms (SNPs) were then called and filtered at the population level using the *populations* module of Stacks. Key pipeline parameters were optimized on a subset of 22 individuals (one per population with the highest and lowest read depth), including *m* (minimum number of identical raw reads for a putative allele), *M* (allowed mismatches between alleles to form a locus), and *n* (allowed mismatches between loci). Following Paris et al., 2017 recommendations, *M* and *n* were varied (fixing *M* = *n* from 1 to 9), while *m* was set to 3 (Paris *et al*., 2017 ; Rochette & Catchen, 2017). The combination of -*m* 3, -*M* 3, and -*n* 3 maximized the number of assembled loci and the level of polymorphism (Appendix B) and was applied to the full dataset. SNPs were retained if genotyped in at least 80% of individuals per population and present in a minimum of 9 out of 11 populations. SNPs with observed heterozygosity >70% or minor allele frequency (MAF) <1% were excluded. The final genotype matrix was exported in VCF format. VCF data were converted into genlight and genind objects using ‘vcfR’ R package (Knaus & GrÜnwald, 2017), and .geno file using ‘LEA’ R package (Frichot & FranÇois, 2015) for downstream analyses.

### Population genomics analyses

Genetic structure was explored using a Principal Component Analysis (PCA) on individual genotypes with ‘adegenet’ R package (Jombart & Ahmed, 2011). Admixture proportions were inferred using sparse non-negative matrix factorization (sNMF) implemented in LEA, testing K = 1–12 with 100 runs per K. The optimal K was chosen from cross-entropy profiles, and individual ancestry coefficients from the best run were visualized with ‘pophelper’ R package (Francis, 2017).

Within-population diversity was assessed through observed and expected heterozygosity (*H*_O_ and *H*_E_), allelic richness (*Ar*), and the inbreeding coefficient (*F*_IS_) computed with ‘hierfstat’ R package (Goudet & Jombart, 2022). Genetic differentiation was quantified using the beta estimator of *F*_ST_ as implemented in ‘hierfstat’ package. This approach provides both population-specific and overall measures of genetic differentiation. Confidence intervals for *F*_IS_ and *F*_ST_ were obtained by bootstrap resampling across loci (10,000 replicates). Private alleles per population were computed using ‘poppr’ R package (Kamvar *et al*., 2014). Between-population genetic differentiation was quantified using Weir and Cockerham’s pairwise *F*_ST_ (WC84) implemented in ‘hierfstat’ (Appendix C). Isolation by distance was tested using a Mantel test (1000 permutations; ‘ade4‘ package (Thioulouse *et al*., 2018)) between geodesic geographic distances (km) and linearized genetic distances (*F*_ST_ / (1 - *F*_ST_); Rousset, 1997). Linear regression was used to visualize the relationship between genetic and geographic distances.

### Species distribution models

To investigate how climate change shaped and could shape the species’ spatial dynamics, we performed species distribution models (SDMs) with both paleo- and future projections, allowing us to formulate hypotheses about its geo-demographic and genetic history and to anticipate its trajectory under climate change. SDMs followed the pipeline detailed in Collette *et al*., (2026), with input variables obtained from the CHELSA dataset, which provide high-resolution (30 arc sec, ∼1 km) layers suitable for paleo (CHELSA-TraCE21k, Karger et al., 2023), present and future (CHELSA 2.1, Karger et al., 2017) bioclimatic and snow cover duration (scd) projections. Soil variables were extracted from SoilGrids 2.0 (Poggio *et al*., 2021) with ‘geodata’ R package (Hijmans, 2025) at 1 km resolution and were assumed to remain constant through time for projections. Correlated exploratory variables (Pearson r > 0.9) were filtered based on their ecological relevance for *X. scabra* (Appendix D). The final set of predictors comprised annual mean temperatures, temperatures of the warmest and coldest quarters, annual precipitation, precipitation of the warmest and coldest quarters, soil depth, coarse fragments, clay, sand and silt content (30–60 cm), and slope. Models were calibrated over the spatial extent encompassing both the Pyrenees and the Alps (longitude: −11, 27 latitude: 34, 60) in order to capture environmental conditions beyond the current distribution of *X. scabra* in its current native range, as some species have historically dispersed from the Alps into this region (Charrier *et al*., 2014 ; SchÖnswetter *et al*., 2002). Models were projected onto major climatic intervals since the Last Glacial Maximum (LGM, 26,500-17,500 years before present (BP)), Heinrich Stadial 1 (17,500-14,700 BP), Bølling-Allerød (14,700-12,900 BP), Younger Dryas (12,900-11,700 BP), and the Early (Greenlandian (11,700-8,200 BP)), Mid (Northgrippian (8,200-4,200 BP)), and Late Holocene (Meghalayan (4,200 BP-1950)). For each interval, projections were based on centennial paleoclimatic time slices from CHELSA-TraCE21k, corresponding to the approximate midpoint of each period (i.e. 20.0, 15.9, 13.8, 12.3, 10.0, 6.2, and 2.1 ka BP). These correspond to CHELSA time indices −200, −159, −138, −123, −100, −62, and −21, respectively. Models were also projected onto present-day conditions (1981-2010), and for future conditions up to 2100 (2011-2040, 2041-2070, 2071-2100, referred to as 2025, 2055 and 2085 in this study) under Shared Socioeconomic Pathways SSP126, SSP370, and SSP585, representing a gradient from optimistic to severe climate scenarios. Response curves generated with the ‘plotmo’ package (Milborrow, 2024) for each environmental variable are shown only for the model with the highest evaluation scores (GAM model, run 6/10, AUC-PR = 0.992, Boyce index = 1, Sensitivity = 1), as it performed best in terms of calibration, discrimination, and classification capacity. Resulting maps were then cropped to the eastern Pyrenees to focus on the study area, while full European projections are provided in Appendix E. Subsequent analyses were performed using Pyrenean binary suitability rasters, obtained by applying a threshold to continuous model predictions, following the approach described by Collette *et al*., 2026.

### Environmental differentiation analyses

To investigate whether genetically differentiated populations also occupied distinct environmental conditions, environmental variables used in species distribution models were extracted for each sampled individual and averaged at the population level. A PCA based on these population-level environmental means was then performed to explore environmental differentiation among populations. Based on the exploratory PCA and SDM response curves, populations were further grouped into “Queralbs” and “Other” categories to test the hypothesis that Queralbs populations occupy distinct environmental conditions. Univariate differences for each environmental variable were assessed using Wilcoxon rank-sum tests with Benjamini–Hochberg correction, while multivariate environmental differentiation between groups was evaluated using PERMANOVA based on Euclidean distances and 10,000 permutations.

### Demographic history analyses

To assess whether climatic and genetic patterns were associated with demographic events, we inferred temporal changes in effective population size (Nₑ) using Stairway Plot v2 (Liu & Fu, 2020). This method estimates demographic history from the site frequency spectrum (SFS) using a flexible multi-epoch model without requiring reference genome or whole-genome sequencing data. Demographic inference was based on a folded SFS generated from the filtered VCF file using *easySFS.py* (https://github.com/isaacovercast/easySFS). Projection values selected from the easySFS preview were 20 for all populations, except Tartera de Ferreroles (8). Stairway Plot analyses were performed with 200 bootstrap replicates and multiple random breakpoints (nrand = 4, 9, 14, and 18 for populations projected to 20 sequences; nrand = 1, 3, 5, and 6 for Tartera de Ferreroles). We assumed a mutation rate of μ = 7 × 10⁻⁹ per site per generation, an estimation of angiosperm autosomal markers (Krasovec *et al*., 2018) and a generation time of 3 years. Outputs were summarized as *N*ₑ trajectories through time with 95% confidence intervals and plotted against years BP, with shaded intervals highlighting the major Late Quaternary climatic periods considered in the SDM analyses.

To complement these SFS-based reconstructions, we performed demographic model inferences using DILS (Demographic Inferences with Linked Selection, FraÏsse *et al*., 2021). Individuals were partitioned into two distinct geographic clusters separated by the Segre Valley (West: Cabaneta, Rialb, Tartera de Ferreroles, Castellet de Moro; East: Mantet, Eyne, Puigmal, Queralbs, Tosa d’Alp, Cadí). DILS was used to compare alternative divergence scenarios based on genomic summary statistics and folded SFSs from noncoding loci. Following the hierarchical framework implemented in DILS, models with current isolation (SI and AM) were first compared to models with ongoing migration (IM and SC), after which genomic models assuming homogeneous versus heterogeneous effective population size and migration rates across loci were evaluated. Analyses were performed under a constant population size model (population_growth = constant) and a bimodal barrier model (modeBarrier = bimodal). Prior ranges (Tsplit: [0–60k] generations; Ne: [500–100k]; M: [1–40]) were informed by Stairway Plot results. The mutation rate was fixed at μ = 7 × 10⁻⁹ and the recombination-to-mutation ratio at ρ/θ = 0.5. Model choice and fit were assessed through posterior predictive checks comparing observed and simulated summary statistics (Goodness-of-Fit).

## Results

### Moderate genetic diversity and limited inbreeding despite small population sizes

After filtering, the dataset retained 21,970 SNPs across 125 individuals from the 11 sampled populations (Figure 1). Overall, the dataset showed relatively low levels of missing data (10.15% on average, Appendix F).

Within-population genetic diversity was moderate (Table 1), with mean expected heterozygosity of 0.165 (*H*_E_ = 0.151 in Queralbs 1 to 0.187 in Cadí) and mean observed heterozygosity of 0.153 (*H*_O_ = 0.139 in Queralbs 2 to 0.173 in Cadí). Inbreeding was close to 0 (*F*_IS_ = 0.074 on average, 0.05 in Cabaneta to 0.1 in Mantet). All *F_IS_* estimates fell within 95% bootstrap confidence intervals. Allelic richness (*Ar*), standardized to the minimum sample size and averaged across variable sites, ranged from 1.311 (Queralbs 1-2) to 1.390 (Cadí) across populations (mean = 1.338), with moderate variation among loci within populations (SD = 0.35-0.41). The number of private alleles (*Ap*) averaged 700 across populations, with Mantet exhibiting the highest number (1,128) and Queralbs 2 the lowest (96). Genetic differentiation was high (overall *F_ST_* = 0.157, 95% CI [0.155; 0.160]) indicating strong population structure. Population-specific *F_ST_* values, estimated relative to the overall genetic structure across all populations, ranged from 0.042 in Cadí to 0.224 in Queralbs 1. Cadí appeared weakly differentiated from the overall population structure, whereas the Queralbs populations were the most genetically differentiated.

**Table 1.**
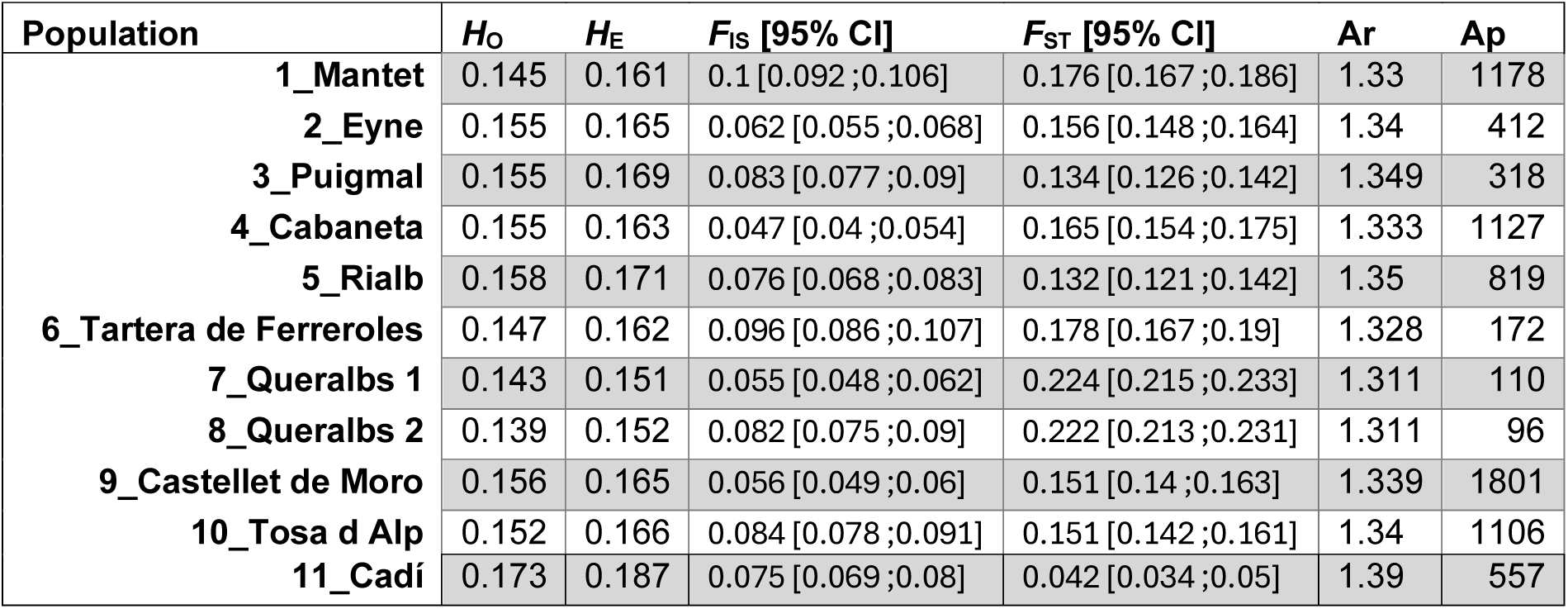
Summary genetic statistics for the 11 populations of *Xatartia scabra*. Statistics include observed heterozygosity (*Ho*), expected heterozygosity (*He*), the inbreeding coefficient (*F_IS_*) with 95% confidence intervals, mean allelic richness (*Ar*) across loci, the number of private alleles (*Ap*), and mean pairwise *F_ST_* with 95% confidence intervals. Mean pairwise *F_ST_* represents the average genetic differentiation of each population relative to all others.

### Genetic structuration among sky-island summits

The PCA revealed a clear separation of populations according to their summit, with the first two axes explaining 12.33% of the total genetic variance (Figure 2A). Populations from geographically distant summits formed more distinct clusters along the first two PCA axes. The significant correlation between genetic and geographic distances (linear regression: R² = 0.489, p = 2.92e-9; Mantel p < 0.001; Figure 2B) indicates isolation by distance.

**Figure 2.**
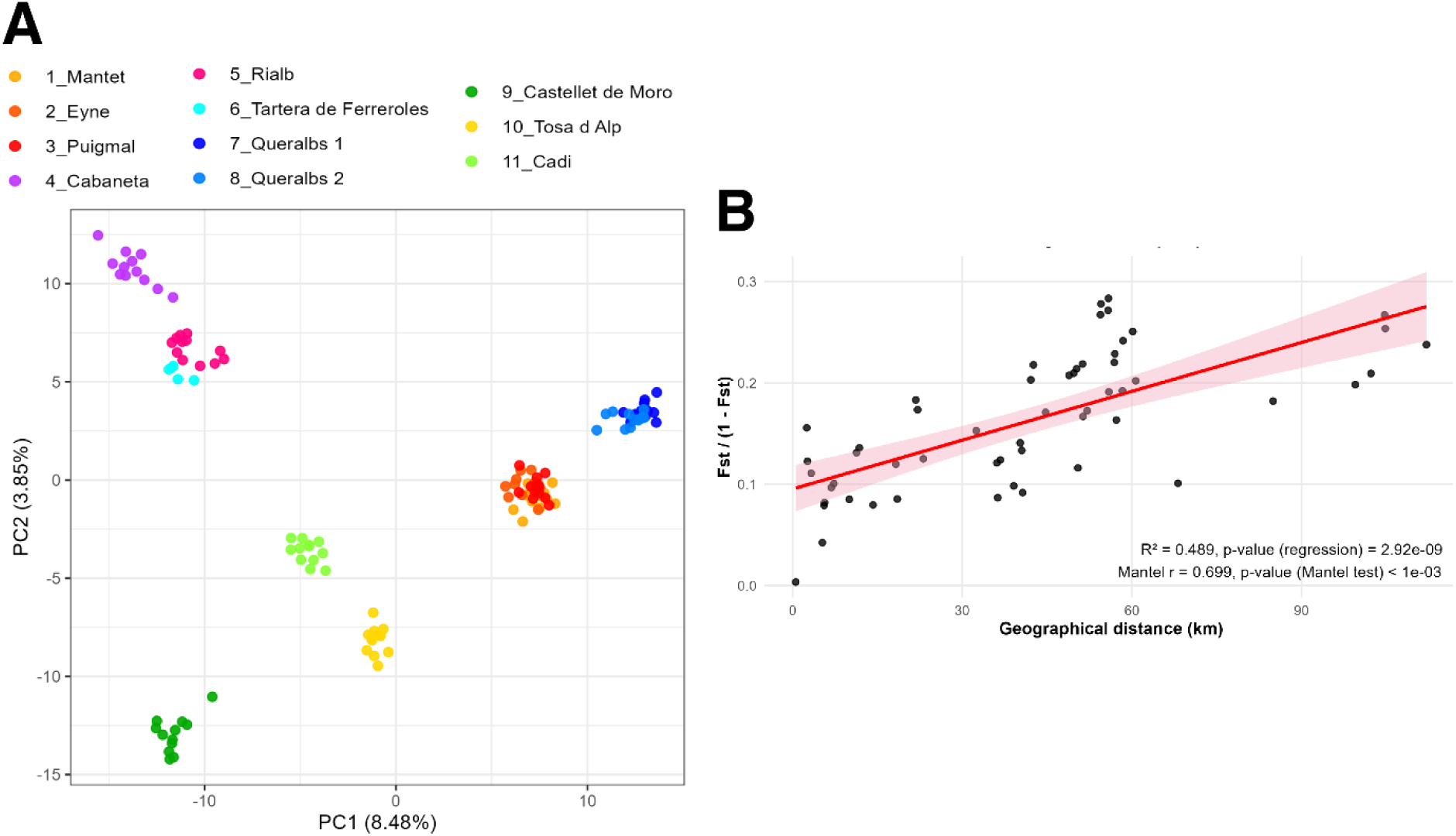
Population genetic structure and isolation-by-distance in *Xatartia scabra.* **(A)** Principal Component Analysis (PCA) based on 21,970 SNPs for the 11 sampled populations of *X. scabra*. Individuals are plotted according to the first two principal components (PC1 and PC2), which explain 8.48% and 3.85 % of the total genetic variance, respectively. Points are colored according to population origin. The Castellet de Moro individual clustering near the center reflects missing data (Appendix F). **(B)** Relationship between pairwise genetic differentiation and geographical distance between populations, showing a significant isolation-by-distance pattern (linear regression R² = 0.489, p < 0.001; Mantel test p < 0.001).

The sNMF analysis supported K = 6 main genetic clusters (Appendix G). These corresponded to Queralbs 1–2 (same summit), the three French populations (same massif), two Andorran populations (Rialb and Tartera de Ferreroles), Cabaneta, Tosa d’Alp, and Castellet de Moro (Figure 3A). French populations showed partial genetic similarity with Queralbs and, for some individuals, with Tosa d’Alp. Within the Andorran group, Tartera de Ferreroles showed genetic similarity with Cabaneta and Rialb, while Castellet de Moro formed a distinct cluster. Cadí showed a mixed genetic profile, sharing ancestry components with all populations, consistent with its lower genetic differentiation from the others (Table 1, Figure 3A).

**Figure 3.**
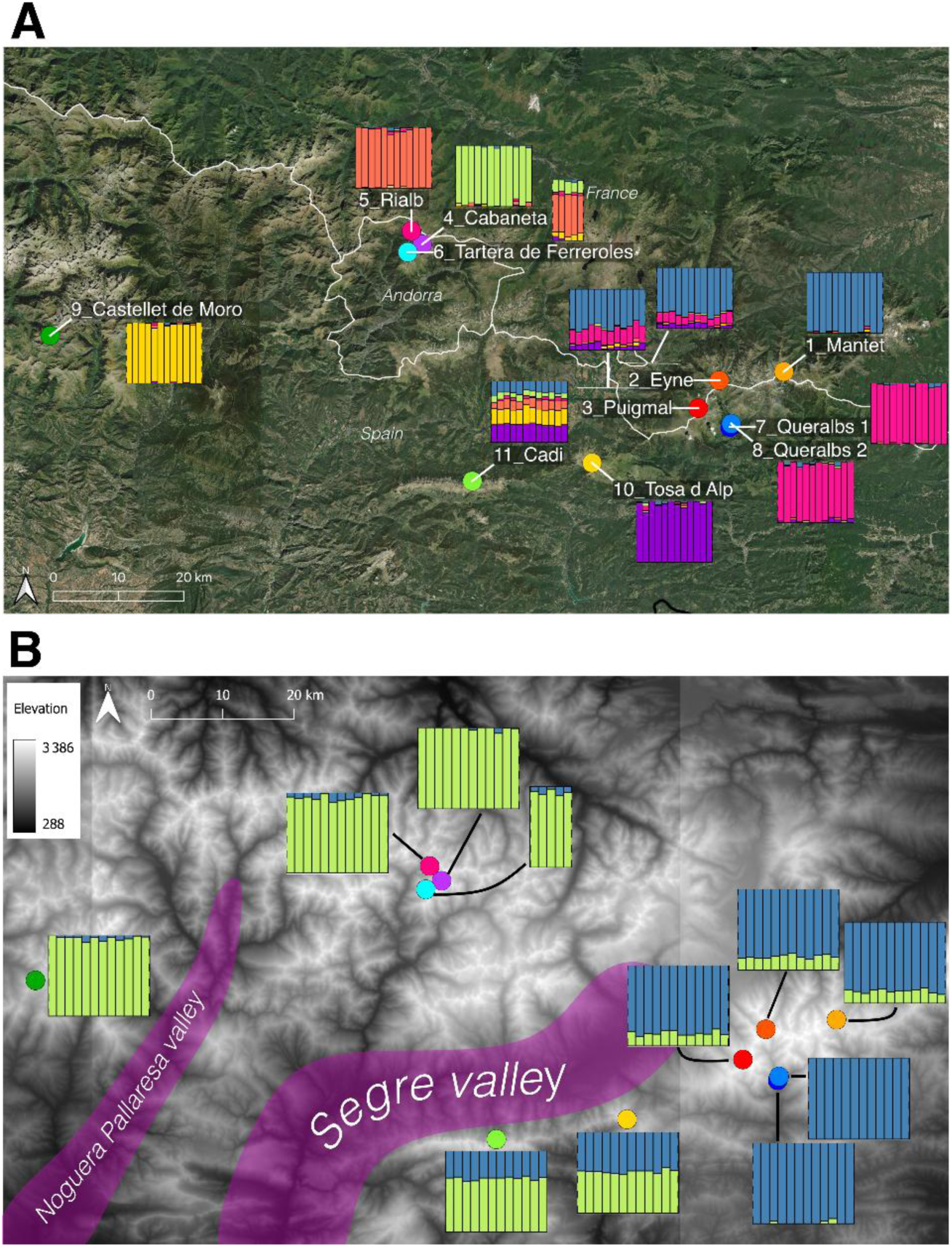
Spatial distribution of genetic ancestry in *Xatartia scabra* inferred with sNMF. Spatial representation of genetic structure across the eastern Pyrenees based on sNMF analyses of 21,970 SNPs from 125 individuals of *X. scabra*. Each barplot corresponds to a sampled locality, with vertical bars representing individuals and colors indicating their estimated ancestry coefficients. **(A)** Genetic structure inferred assuming K = 6 ancestral populations. Barplots are positioned at the geographic location of each sampled population, illustrating the spatial distribution of ancestry components across the study area. **(B)** Genetic structure inferred assuming K = 2 ancestral populations. This configuration highlights a major east–west genetic split among populations. The Segre Valley is indicated as a potential geographic barrier to gene flow separating the two genetic clusters. Sampling locations are marked by colored dots, and elevation is represented by the grayscale background.

Interestingly, K = 2 (Figure 3B, Appendix G) indicated a pronounced east–west genetic partition, consistent with the species’ geographic distribution and the barrier effect of the Segre Valley separating eastern from western populations. This spatial structure was thus used to define population groups and test alternative divergence scenarios with DILS.

### Decline in effective population size since the Last Glacial Maximum

Demographic reconstructions showed relatively high estimated Nₑ during the Late Quaternary, followed by a decline starting after the Last Glacial Maximum (Figure 4). Nₑ decreased from ∼80,000 individuals to current values estimated between 700 and 9,000, although confidence intervals remain broad. Current estimates of Nₑ are higher than observed census sizes (at most a few hundred). The demographic decline coincides with climate, consistent with the onset of population isolation.

**Figure 4.**
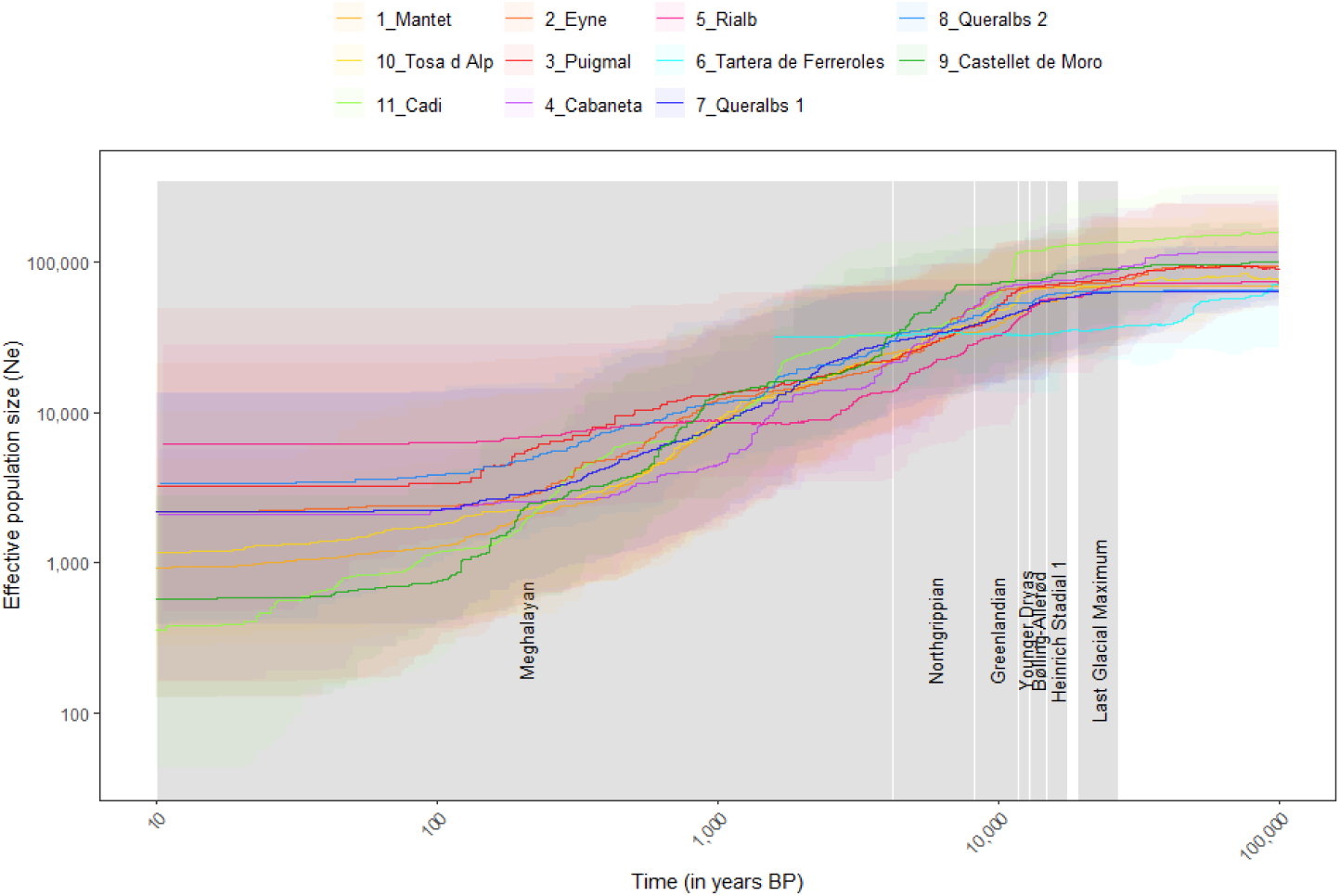
Demographic history of *Xatartia scabra* populations inferred with Stairway Plot. Changes in effective population size (Nₑ) through time for the 11 studied populations of *X. scabra*, inferred using Stairway plot v2 from folded site frequency spectra derived from 21,970 SNPs. Colored lines represent the mean estimates of Nₑ for each population, and shaded areas indicate the corresponding 95% confidence intervals. Major late Quaternary climatic periods are indicated in the background for reference: Last Glacial Maximum (26,500-17,500 years BP), Heinrich Stadial 1 (17,500-14,700 BP), Bølling-Allerød (14,700-12,900 BP), Younger Dryas (12,900-11,700 BP), and the three Holocene stages: Greenlandian (11,700–8,200 BP), Northgrippian (8,200–4,200 BP), and Meghalayan (4,200 BP–1950 CE).

Demographic modeling supported scenarios involving gene flow, although the data did not clearly distinguish between isolation-with-migration and secondary contact. The ancestral Nₑ was estimated between ∼60,000 and 110,000 individuals, followed by a reduction after divergence. However, divergence time was highly sensitive to prior settings. Estimated divergence times varied broadly across runs, corresponding to approximately 100,000 to 400,000 years ago (assuming 3 years per generation). The relative effective population sizes of the East and West clusters were not consistent across runs (Appendix H).

### From past connectivity to future contraction under climate change

Paleoclimatic projections using the current baseline (1981–2010) as a reference indicate that abiotic suitable area was consistently more extensive in the past than today (Figure 5, Figure 6), with higher values during the LGM (+19.6%), Bølling–Allerød (+13.8%), early Holocene (+9.2%), mid-Holocene (+14.9%) and late Holocene (+33.9%), and maxima during Heinrich Stadial 1 (+58.4%) and the Younger Dryas (+41.2%). These results suggest that suitable habitats were generally more widespread across the eastern Pyrenees during the Late Quaternary, with particularly high connectivity during cold post-LGM phases.

**Figure 5.**
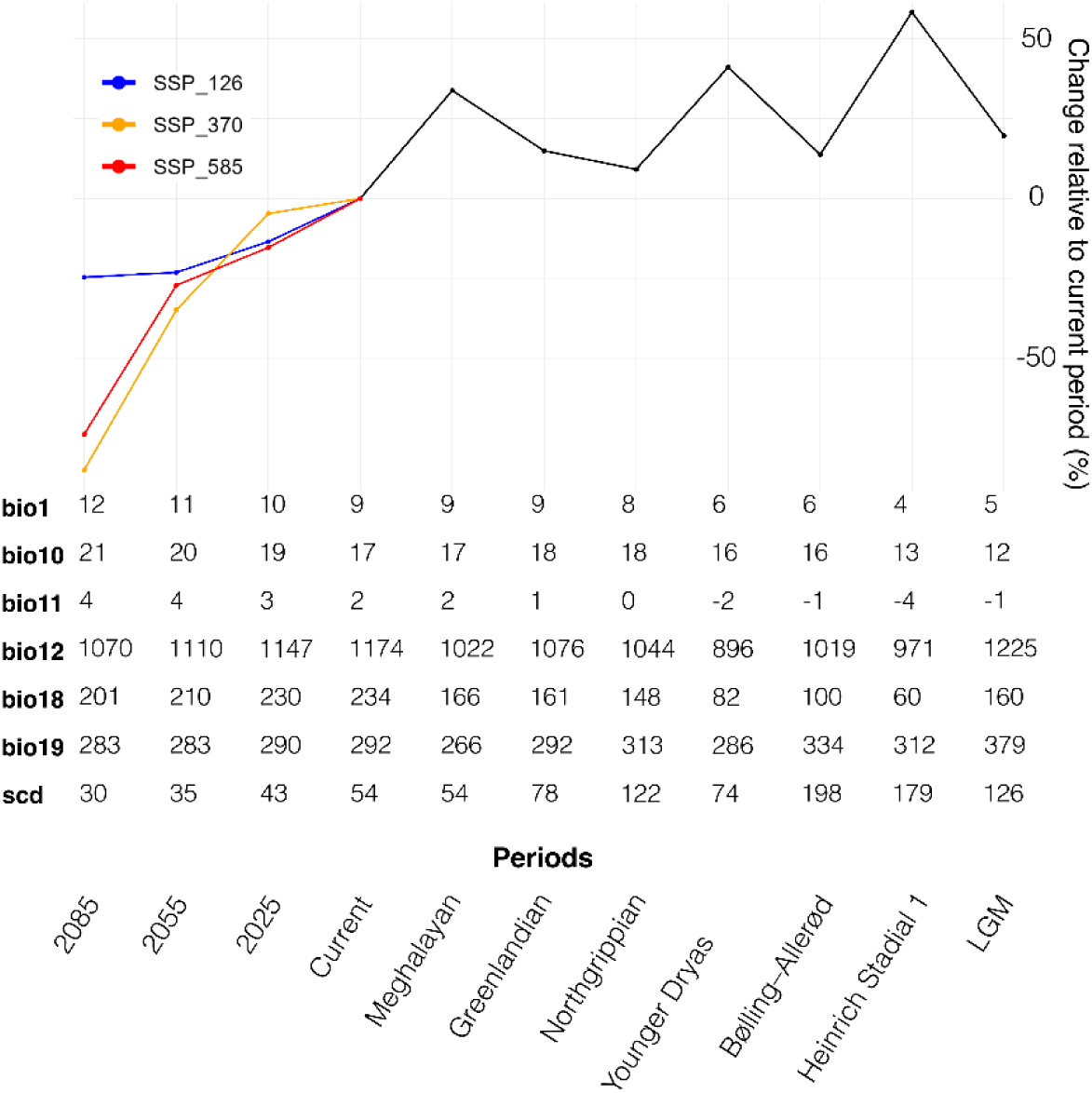
Temporal changes in abiotic suitability for *Xatartia scabra* from the Last Glacial Maximum to future climate scenarios. Values represent the relative change (%) in suitable areas compared with the current baseline climate (1981–2010). Past periods (black line) include the Last Glacial Maximum (26,500–17,500 years BP), Heinrich Stadial 1 (17,500–14,700 BP), Bølling–Allerød (14,700–12,900 BP), Younger Dryas (12,900–11,700 BP), and the three Holocene stages: Greenlandian (11,700–8,200 BP), Northgrippian (8,200–4,200 BP), and Meghalayan (4,200 BP–1950 CE). Future projections (colored lines) correspond to three Shared Socioeconomic Pathways (SSP126, SSP370, SSP585) for the periods 2011-2040, 2041-2070, 2071-2100 (referred to as 2025, 2055 and 2085). Values are shown only for the environmental predictors retained in the model: Bio1 (Annual Mean Temperature, °C), Bio10 (Mean Temperature of Warmest Quarter, °C), Bio11 (Mean Temperature of Coldest Quarter, °C), Bio12 (Annual Precipitation, mm), Bio18 (Precipitation of Warmest Quarter, mm), Bio19 (Precipitation of Coldest Quarter, mm), and SCD (Snow Cover Duration, days).

**Figure 6.**
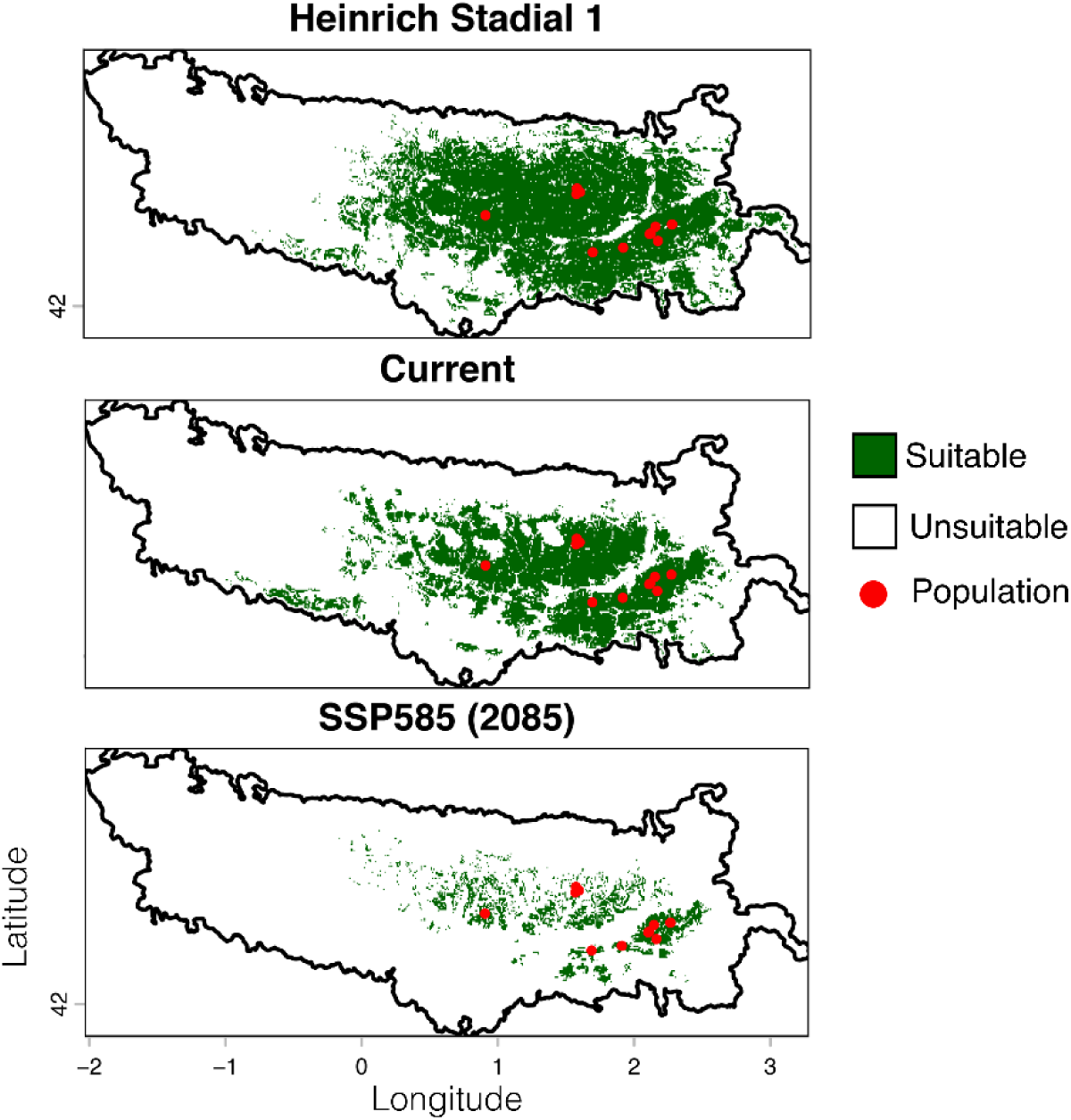
Changes in the spatial distribution of bioclimatic suitable areas for *Xatartia scabra*. Binary projections of bioclimatic suitable areas for *X. scabra* within the Pyrenees for three time periods: Heinrich Stadial 1 (17,500–14,700 BP), the current climate (1981–2010), and a future projection under the SSP585 scenario for 2071-2100 period. Green pixels indicate bioclimatic suitable areas, and white pixels indicate unsuitable areas based on the binarized species distribution models. Red dots represent the species’ current distribution.

Future projections, relative to the current climate (1981–2010), indicated range contractions under climate change (Figure 5 and Figure 6). By 2071–2100, suitable habitat is projected to decline by −24.7% under the most optimistic scenario (SSP126), −85.0% under SSP370, and −73.9% under the high-emission scenario SSP585 (Figure 6), representing an average reduction of −61.2% across scenarios.

These changes have occurred in the context of climatic shifts since the Late Quaternary. Mean annual temperature across the Pyrenees increased from approximately 4 °C during Heinrich Stadial 1 (17,500-14,700 BP) to about 8 °C under current conditions (1981–2010) and is projected to reach 12 °C by 2080-2100 period on average across SSPs scenarios (Figure 5). Over the same period, the snow cover duration declined from 179 days during Heinrich Stadial 1 to 54 days today and is projected to decrease further to 30 days by 2085. Fluctuations in suitable area extent from the LGM to the present (Figure 5) appeared to track temporal variation in both precipitation- and temperature-related variables, with range expansions generally associated with colder and drier climatic conditions.

### Southeastern populations at the edge of the species’ climatic niche

Response curves from the calibration model and the PCA of environmental variables (Figure 7A, Figure 7B), in which the first two axes explained 83.4% of the total environmental variation among populations (PC1 = 67.8%, PC2 = 15.6%), suggested that the two Queralbs populations (Queralbs 1 and Queralbs 2) occupy distinct environmental conditions compared with the other populations. Queralbs populations were associated with higher temperatures, including annual mean temperature and temperatures of the warmest and coldest quarters (+ 5.7 °C), as well as substantially lower snow cover duration (- 130 days). Univariate analyses revealed similar trends for several climatic and soil variables (Figure 7), although none remained statistically significant after Benjamini–Hochberg correction. In contrast, PERMANOVA detected significant multivariate environmental differentiation between Queralbs and the remaining populations (R² = 0.497, p = 0.019), indicating that nearly 50% of the environmental variance was associated with group identity.

**Figure 7.**
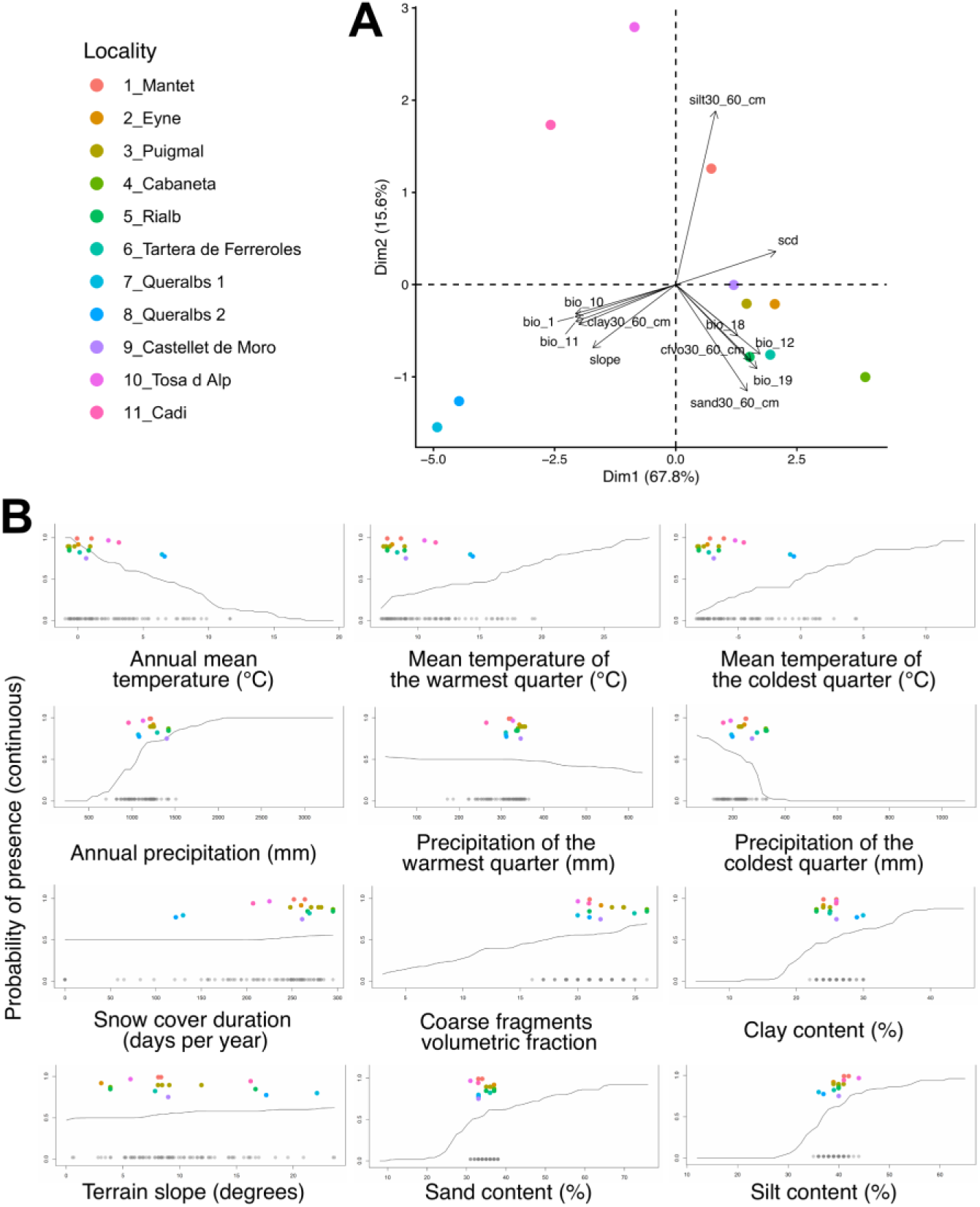
Environmental differentiation analyses for *Xatartia scabra*. **(A)** Principal component analysis (PCA) of environmental conditions across the 11 populations of *Xatartia scabra*. Arrows indicate the contribution of climatic, soil, and topographic variables to the ordination space. The first two axes explained 67.8% (PC1) and 15.6% (PC2) of the total environmental variance, respectively. Climatic variables included annual mean temperature (bio_1), temperatures of the warmest (bio_10) and coldest (bio_11) quarters, annual precipitation (bio_12), precipitation of the warmest (bio_18) and coldest (bio_19) quarters, and snow cover duration (scd). Soil variables included coarse fragment volumetric content (cfvo30_60_cm), clay, sand, and silt contents at 30–60 cm depth. **(B)** Response curves of the species distribution model are shown only for the best-performing model (GAM model, run 6/10; AUC-PR = 0.992, Boyce index = 1, Sensitivity = 1). Black lines represent partial dependence curves showing the average effect of each predictor on the predicted probability of presence while averaging over the observed values of the other predictors. Gray points indicate the distribution of presence records used for model calibration. Colored points correspond to the environmental conditions at the sampling locations of the 11 studied populations; their vertical position is arbitrary to avoid overlap and does not represent predicted probabilities.

## Discussion

In this study, we tested whether the genetic patterns of *X. scabra* match the expectations of a classical sky island system, in which populations are restricted to isolated summits and gene flow is strongly limited by topography. Our results largely support this framework, with pronounced genetic structuring and a significant isolation-by-distance pattern, consistent with limited gene flow among populations. However, not all predictions were met. Despite strong spatial isolation and small census sizes, inbreeding levels remained low across populations, and effective population size estimates were higher than expected given the inferred demographic decline and field observations. This apparent mismatch suggests that current fragmentation has not (yet?) resulted in the level of genetic erosion typically predicted under such conditions. It further indicates that biological processes, such as life-history traits or buffering mechanisms maintaining diversity, must be considered to interpret these patterns. Together, these results provide a nuanced view of how isolation, demography, and species-specific traits interact to shape genetic variation in a sky island context.

### Topographic barriers and glacial connectivity shape the genetic structure of *Xatartia scabra*

While present-day patterns clearly reflect strong spatial isolation among summit populations, paleoclimatic projections suggest that connectivity is likely to have been higher during cold Late Quaternary phases. During Heinrich Stadial 1 (17,500-14,700 BP), suitable climatic conditions likely extended into lower-elevation areas, potentially allowing intermittent connectivity among summits that are currently isolated. Under such conditions, valleys may have acted less as barriers and more as corridors facilitating gene flow. Conversely, during warmer periods following the Last Glacial Maximum (26,500-17,500 BP), the upward contraction of suitable habitats would have progressively restricted populations to high-elevation summits. In the conical topography of the Pyrenees, where available areas decrease with elevation, this process likely reinforced fragmentation, contributing to the strong genetic structure observed today.

At finer spatial scales, admixture patterns likely reflect geographic proximity and relatively recent divergence among neighboring summits, suggesting that gene flow has primarily occurred among nearby populations, potentially along ridgelines. In contrast, deep valleys, such as the Segre Valley, appear to act as recurrent barriers structuring genetic variation in the eastern Pyrenees, as reported in other taxa (Bech *et al*., 2009 ; 2013 ; Buso *et al*., 2026 ; Valbuena-UreÑa *et al*., 2018). Castellet de Moro is further isolated by the Noguera Pallaresa Valley, reinforcing the isolation for this population. This spatial structure is likely strengthened by the dispersal ecology of *X. scabra*, as anemochorous seed dispersal may be constrained by topography, limiting effective cross-valley exchange. Similar patterns of summit-scale isolation have been reported in other alpine organisms, highlighting the role of topographic barriers in shaping genetic structure (Hartley *et al*., 2023 ; Morente-Lopez *et al*., 2018 ; Polato *et al*., 2017 ; Salvado *et al*., 2022 ; Slatyer *et al*., 2014).

Within this framework, the Cadí mountain range stands out as a particularly informative case. Its relatively low genetic differentiation, high genetic diversity, and admixed ancestry profile are consistent with a potential role as a connectivity hub during colder periods. Its central geographic position further supports this interpretation. However, alternative explanations must be considered. The observed genetic signature may also reflect the persistence of ancestral variation if Cadí corresponds to an ancestral population or a refugium, rather than a simple zone of secondary contact. Distinguishing among these scenarios remains challenging, especially in the absence of genome-wide data, and similar patterns reported in another Pyrenean endemic (Salvado *et al*., 2022) suggest that additional processes may be involved.

Demographic reconstructions indicate a sustained postglacial decline in Nₑ since the Last Glacial Maximum, consistent with habitat contraction and the progressive loss of connectivity inferred from paleoclimatic projections, a pattern also reported in several plant taxa (Deng *et al*., 2026 ; Han *et al*., 2025 ; Vintsek *et al*., 2022). However, the observation that present-day Nₑ estimates exceed census sizes (N) across populations is unexpected (Ellstrand & Elam, 1993 ; Waples, 2025). This discrepancy may partly reflect methodological and technical biases, as SFS-based inference can be sensitive to limited sample sizes (e.g. excess low-frequency variants due to PCR duplicates; <25,000 SNPs; <15 individuals; Nunziata & Weisrock, 2018). At the same time, such patterns may also arise from biological processes that decouple effective from census size, such as persistent seed banks that can increase long-term effective population size by buffering genetic drift across generations. Distinguishing between these alternatives remains challenging, but this ambiguity raises the possibility that genetic diversity may be more effectively maintained than expected under strong isolation. Our demographic results point to a divergence history shaped by gene flow, but key parameters remain weakly constrained. The broad range of divergence time estimates (∼100,000–400,000 years ago) spans several Pleistocene climatic cycles, from the onset of the last glacial period to earlier glacial–interglacial phases. However, the strong sensitivity of the inference to prior settings prevents discrimination between these alternative temporal scenarios and limits conclusions on demographic asymmetries between clusters. These patterns may also reflect the influence of species-specific life-history traits, although this cannot be directly assessed with the current data.

### Genetic resilience persists despite isolation, yet adaptive potential remains uncertain

Despite strong postglacial fragmentation, long-term isolation among summits, small census sizes and restricted distribution, *X. scabra* retains a non-negligible level of genetic diversity. Inbreeding coefficients remained consistently low across all populations, contrasting with the elevated inbreeding commonly reported in narrow-ranged mountain taxa (Salvado *et al*., 2022 ; 2026; but see ÆgisdÓttir *et al*., 2009). In this regard, *F*_IS_ values measured in this study are similar to the ones found with the same genotyping approach (nGBS) on strictly allogamous orchids whose seeds can be wind-dispersed across long distances (Gibert *et al*., 2024 ; Salvado *et al*., 2025). This pattern suggests that substantial genetic variation has been maintained, with no clear evidence of severe diversity erosion to date, although genetic erosion may occur with a temporal lag following demographic decline (Aguilar *et al*., 2009 ; Gargiulo *et al*., 2025 ; Pinto *et al*., 2024). In this context, the unexpectedly high Nₑ estimates may not solely reflect methodological artifacts but could also point to biological mechanisms buffering genetic diversity.

Several non-exclusive processes may explain this apparent resilience. First, limited gene flow among nearby populations may persist, likely mediated by pollinators, and contribute to maintaining genetic diversity despite overall differentiation (Smith & Pauli, 2024). Second, the absence of strong inbreeding is consistent with a strictly sexual, obligately outcrossing system, potentially reinforced by self-incompatibility. In such systems, negative frequency-dependent selection at the S-locus maintains allelic diversity and promotes outcrossing, helping preserve genome-wide variation even in fragmented populations (Barrett, 2003 ; Pickup & Young, 2008 ; Silva *et al*., 2016). In addition, the monocarpic life history of *X. scabra* may further contribute to maintaining genetic diversity. By concentrating reproductive effort on a single event, monocarpic species may reduce variance in individual reproductive success and allow a large proportion of individuals to contribute to the next generation, potentially increasing effective population size relative to census size. When combined with obligate outcrossing and overlapping generations, typical of long-lived alpine perennials, this life-history strategy may further limit inbreeding (ÆgisdÓttir *ET AL.*, 2009). Consistent with this interpretation, strong inbreeding depression may reduce the persistence of selfed or closely related offspring, as inbred individuals often exhibit reduced fitness and may be selectively eliminated before reaching reproductive stages, resulting in low observed inbreeding coefficients even in small populations (Keller & Waller, 2002). Finally, seed banks may also buffer genetic diversity by storing alleles across generations and slowing the loss of variation through genetic drift, particularly under fluctuating demographic conditions (McCue & Holtsford, 1998).

More broadly, these results raise the question of whether high levels of genetic diversity and relatively large effective population sizes necessarily translate into adaptive potential under ongoing environmental change or whether will be able to track rapidly shifting climates through range shifts (Oliveira *et al*., 2026). However, the maintenance of neutral genetic diversity does not necessarily imply that populations retain sufficient adaptive variation to cope with ongoing environmental changes, and whether the standing genetic variation observed includes adaptive alleles remains unknown. In this context, environmental predictors derived from SDMs indicate that the two Queralbs populations occur at the warm margin of the species’ climatic niche, suggesting possible ecological differentiation. While such marginal populations may experience stronger selective pressures and thus harbor local adaptation contributing to their persistence (Cross & Eckert, 2024 ; Forester *et al*., 2022), they may also be particularly vulnerable to rapid environmental change (Anderson *et al*., 2025). Testing these hypotheses will require genome-wide data capable of detecting finer adaptive signals (Bernatchez *et al*., 2024). Species facing the highest conservation risks or suffering from major data deficiencies tend to remain underrepresented in genomic resources. (Monchamp *et al*., 2023). *X. scabra* exemplifies this gap. Although our data provide valuable insights into patterns of population structure and demographic history, they have limited power to detect loci under selection or genotype–environment associations (LOWRY *et al*., 2017). Linking adaptive genomic variation to fine-scale environmental gradients will be critical to assess the species’ capacity to persist under rapidly changing conditions. Overall, while *X. scabra* shows greater genetic resilience than expected for such a narrowly distributed and fragmented species, its long-term persistence remains uncertain under the combined pressures of climate warming and human disturbance.

### Climate change threatens the future persistence of *Xatartia scabra*

Future climate projections predict a drastic reduction (−61.2 % on average) in suitable abiotic conditions for *X. scabra* by 2071-2100 period, with suitable areas increasingly restricted to the highest ridges. Such contractions mirror patterns reported for many narrow-ranged mountain species, which are expected to lose most of their bioclimatically suitable area under climate change (Collette *et al*., 2026 ; Dullinger *et al*., 2012) and shift upslope (Pauli *et al*., 2012).

These estimates, however, require cautious interpretation. Scree systems are geomorphically dynamic, where snow cover, freeze–thaw cycles, and drought continuously reshaping fine-scale niches beyond the resolution of coarse climatic predictors (Huc, 2010). As a result, such models may overestimate broad-scale climatic suitability while failing to capture microrefugia that can buffer populations and sustain site-level persistence (Chauvier *et al*., 2022 ; Trivedi *et al*., 2008).

Beyond climate, persistence ultimately depends on the availability of suitable microsites within mobile scree. Under current conditions, recruitment is largely confined to small, relatively stable patches where seeds can lodge (Huc, 2010); high juvenile attrition and strong intraspecific competition among adults yield naturally sparse stands (Aymerich, 2003). Climate warming is likely to intensify these constraints as the treeline shifts upslope (Ameztegui *et al*., 2016 ; Delpouve *et al*., 2025), promoting shrub encroachment, already facilitated by ongoing warming (Francon *et al*., 2022), and the progressive stabilization of open scree. In some localities, the expansion of shrubs (e.g. *Rhododendron*) already indicates partial stabilization and a contraction of open, mobile microhabitats, a trend likely reinforced by declining snow cover and increasing progressive drying (Huc, 2010). This trend risks reducing the availability of open, mobile microhabitats on which *X. scabra* depends. At the population scale, flowering now peaks in mid-late July, earlier than historical records (H_UC_, 2010 ; Laurents *et al*., 2024), consistent with ongoing warming (OPCC, 2018) and raising the prospect of plant-pollinator mismatch (Gérard *et al.*, 2020 ; Li *et al*., 2024); no published study has documented the pollinators of this species to date. Earlier flowering could also expose reproductive structures to damage from late frosts (Inouye, 2008). In addition to the documented shift toward earlier flowering, historical accounts portray the species as rare contrast with its current local abundances, indicating that populations have already responded to recent climate change. These signals underscore that demographic processes and microhabitat dynamics must be assessed alongside climate projections: the long-term persistence of *X. scabra* will depend not only on climatic suitability but also on the availability and stability of inherently fragmented scree microhabitats and associated pollinator communities, which are highly vulnerable to disturbance. In addition to climatic threats, *X. scabra* could face increasing anthropogenic pressures in high-mountain environments (Bowler *et al*., 2020 ; Qazi *et al*., 2025). Pastoralism, together with the rising pressure from hiking, especially where scree slopes are crossed off trail (Onete & Pop, 2010), can compromise the stability of the microhabitats on which the species depends, thereby altering vegetation dynamics. Available evidence indicates that natural ungulate grazing (e.g. Pyrenean chamois) is not the primary constraint on viability (Aymerich, 2003), but ovine trampling can locally damage individual plants (Baudiere & and Serve, 1980). These combined pressures are likely to exacerbate fragmentation and isolation, thereby increasing extinction risk (RamÍrez-Delgado *et al*., 2022). Yet their ecological and demographic consequences remain largely undocumented. The combination of ecological specialization, strong habitat dependence, and limited dispersal capacity underscores the heightened vulnerability of this narrow endemic.

### Conservation priorities for a fragmented Pyrenean endemic

From a conservation standpoint, safeguarding existing populations of *X. scabra* remains the most effective strategy to ensure the species’ long-term persistence. Given the species’ restricted range, small census sizes, and pronounced genetic structuring, each population represents a distinct reservoir of genetic and evolutionary history. Maintaining the current mosaic of populations will maximize the preservation of genetic variation and adaptive potential, strengthening the species’ chances of persistence under accelerating climate change (Bowgen *et al*., 2022 ; Hoelzel *et al*., 2019). Within this framework, the Cadí population warrants particular attention, as it appears to play a central role in the species’ evolutionary history. Similarly, the Queralbs populations, located at the warm margin of the species’ range, are of particular interest, as they may exhibit either local adaptation or increased susceptibility to decline, and should therefore be more thoroughly documented.

Conservation actions should prioritize the strict protection of all known sites and the prevention of habitat degradation from tourism, grazing, and infrastructure expansion (Pulido-Chadid *et al*., 2025). Because the species relies on dynamic scree, management should maintain the geomorphological processes sustaining suitable microhabitats, with targeted interventions where shrub encroachment stabilizes scree (Stem *et al*., 2005). Finally, standardized monitoring of population size, recruitment, and habitat conditions is essential for early warning of declines (Jones *et al*., 2013). Annual surveys of the Mantet population spanning more than 30 years provide a ready-to-deploy framework for extending standardized monitoring across populations.(Laurents *et al*., 2024). Coordinated, transboundary management will further enhance these efforts by ensuring consistent protection across the species’ entire range.

Based on current evidence, assisted migration is not recommended to restore gene flow at this stage. While maintaining genetic exchange is important (Turnock *et al*., 2024), natural gene flow remains limited between populations and, if artificially increased, may also carry risks of maladaptive introgression or hybridization (Ellstrand & Elam, 1993). Active interventions can be beneficial yet risk unintended evolutionary outcomes and should therefore be applied cautiously and adaptively (Aitken & Whitlock, 2013 ; GaitÁn-Espitia & Hobday, 2021 ; McKay *et al*., 2005). *In situ* measures can be complemented by *ex situ* safeguards (seed banking, living collections); existing holdings should be increased with material from Cadí to provide insurance against local extinctions. However, technical difficulties and the limited experience with narrowly endemic species highlight the need for caution when applying such approaches (Coelho *et al*., 2020).

## Author Contributions

Conceptualization: NC, JB, VH; Methodology: NC JB PS SP VH; Software: NC JB; Validation: NC JB, VH, SP; Formal analysis: NC; Investigation: NC; Resources: MM JP XO CP GGP IF AG JB; Data curation: NC; Writing – Original Draft: NC, JB; Writing – Review & Editing: NC JB VH SP; Visualization: NC JB; Supervision: JB VH SP; Project administration: JB VH SP; Funding acquisition: VH JB SP.

## Data Archiving

The data for this study have been deposited in the European Nucleotide Archive (ENA) at EMBL-EBI under accession number PRJEB113243 (https://www.ebi.ac.uk/ena/browser/view/PRJEB113243).

