## Supplemental data for "Life-history traits may buffer genetic erosion under isolation in mountain sky-island systems"

**Table of Contents:**

| **Appendix A: Sampling localities, geographic coordinates, and sample sizes of the 11 studied populations of *Xatartia scabra* in the Pyrenees.** | **Page 2** |
| --- | --- |
| **Appendix B: Variation in assembled and polymorphic loci across Stacks parameter settings** | **Page 3** |
| **Appendix C: Pairwise genetic differentiation among the 11 populations of *Xatartia scabra* estimated using Weir and Cockerham’s FST (WC84).** | **Page 4** |
| **Appendix D: Environmental predictors analysis for species distribution modeling** | **Page 12** |
| **Appendix E: European projections of past, current and future abiotic niche suitability continuous maps for *Xatartia scabra*** | **Page 48** |
| **Appendix F: Cross-entropy criterion as a function of the number of ancestral populations** | **Page 52** |
| **Appendix G: Missing data rate per individual (%)** | **Page 53** |
| **Appendix H: DILS model fit and parameter estimates** | **Page 54** |

**Appendix A: Sampling localities, geographic coordinates, and sample sizes of the 11 studied populations of *Xatartia scabra* in the Pyrenees.** Coordinates are given in decimal degrees and correspond to the centroid of each sampled locality.

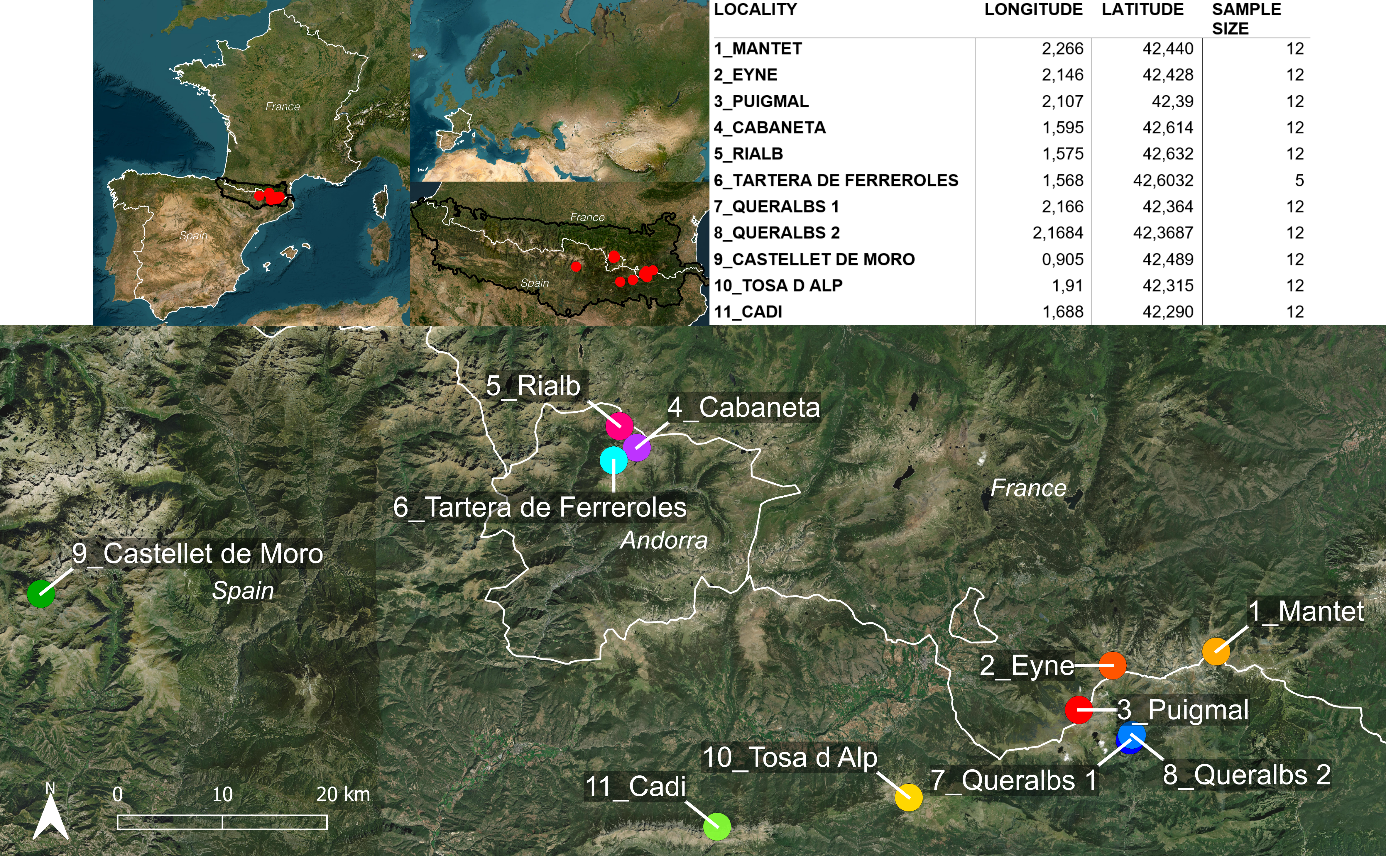

**Appendix B: Variation in assembled and polymorphic loci across Stacks parameter settings**

The number of assembled (dark blue) and polymorphic (light blue) loci obtained under different Stacks parameter combinations (m = minimum number of identical raw reads for a putative allele, M = allowed mismatches between alleles to form a locus, n = allowed mismatches between loci). Key parameters were optimized on a subset of 22 individuals (one per population with the highest and lowest read depth). While *M* and *n* were jointly varied (M = n from 1 to 9) with *m* fixed at 3, the combination *-m 3, -M 3 -n 3* maximized SNP recovery and polymorphic locus assembly and was retained for subsequent analyses.

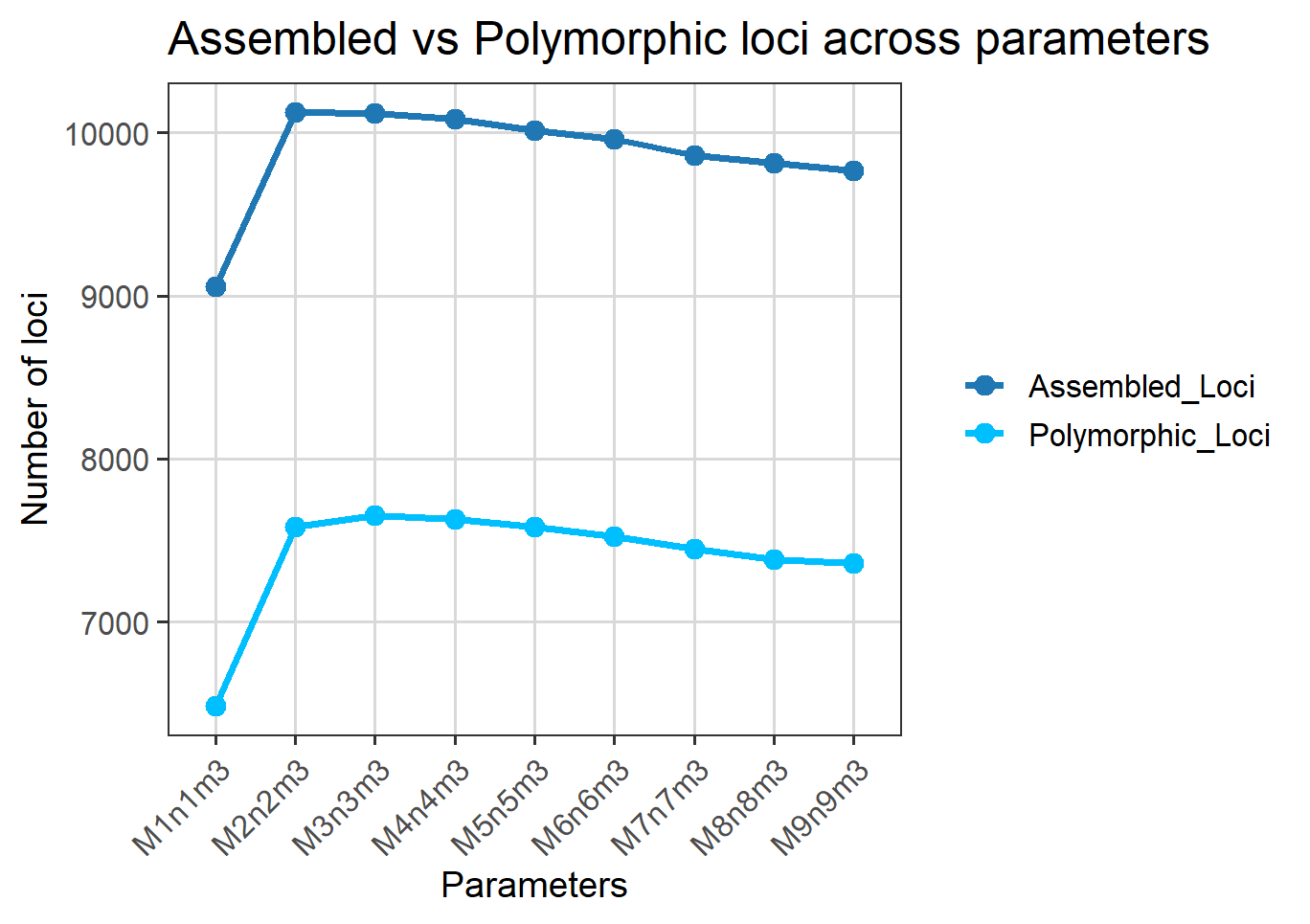

**Appendix C: Pairwise genetic differentiation among the 11 populations of *Xatartia scabra* estimated using Weir and Cockerham’s FST (WC84).**

| **1_Mantet** | **2_Eyne** | **3_Puigmal** | **4_Cabaneta** | **5_Rialb** | **6_Tartera de Ferreroles** | **7_Queralbs 1** | **8_Queralbs 2** | **9_Castellet de Moro** | **10_Tosa d Alp** | **11_Cadi** |  |
| --- | --- | --- | --- | --- | --- | --- | --- | --- | --- | --- | --- |
| NA | 0,078 | 0,074 | 0,195 | 0,168 | 0,201 | 0,120 | 0,116 | 0,192 | 0,133 | 0,104 | **1_Mantet** |
|  | NA | 0,041 | 0,173 | 0,147 | 0,180 | 0,092 | 0,088 | 0,173 | 0,111 | 0,084 | **2_Eyne** |
|  |  | NA | 0,172 | 0,143 | 0,176 | 0,076 | 0,073 | 0,166 | 0,107 | 0,080 | **3_Puigmal** |
|  |  |  | NA | 0,109 | 0,135 | 0,218 | 0,211 | 0,161 | 0,169 | 0,110 | **4_Cabaneta** |
|  |  |  |  | NA | 0,100 | 0,186 | 0,181 | 0,140 | 0,146 | 0,090 | **5_Rialb** |
|  |  |  |  |  | NA | 0,221 | 0,214 | 0,161 | 0,179 | 0,108 | **6_Tartera de Ferreroles** |
|  |  |  |  |  |  | NA | 0,003 | 0,211 | 0,155 | 0,123 | **7_Queralbs 1** |
|  |  |  |  |  |  |  | NA | 0,202 | 0,148 | 0,118 | **8_Queralbs 2** |
|  |  |  |  |  |  |  |  | NA | 0,154 | 0,092 | **9_Castellet de Moro** |
|  |  |  |  |  |  |  |  |  | NA | 0,079 | **10_Tosa d Alp** |
|  |  |  |  |  |  |  |  |  |  | NA | **11_Cadi** |

**Appendix D: Environmental predictors analysis for species distribution modeling**

**Table D1. Description of environmental predictors used in species distribution model.** Bioclimatic variables and snow cover duration were obtained from the CHELSA dataset (30 arc sec resolution; Karger et al., 2017, 2023). Elevation data were derived from the GMTED2010 digital elevation model, and soil variables from SoilGrids v2.0 (Poggio *et al.*, 2021).

| Variable | Description | Unit |
| --- | --- | --- |
| bio_1 | Mean annual temperature | °C |
| bio_2 | Mean diurnal range | °C |
| bio_3 | Isothermality | % |
| bio_4 | Temperature seasonality | °C |
| bio_5 | Max temperature of warmest month | °C |
| bio_6 | Min temperature of coldest month | °C |
| bio_7 | Temperature annual range | °C |
| bio_8 | Mean temperature of wettest quarter | °C |
| bio_9 | Mean temperature of driest quarter | °C |
| bio_10 | Mean temperature of warmest quarter | °C |
| bio_11 | Mean temperature of coldest quarter | °C |
| bio_12 | Annual precipitation | mm |
| bio_13 | Precipitation of wettest month | mm |
| bio_14 | Precipitation of driest month | mm |
| bio_15 | Precipitation seasonality | % |
| bio_16 | Precipitation of wettest quarter | mm |
| bio_17 | Precipitation of driest quarter | mm |
| bio_18 | Precipitation of warmest quarter | mm |
| bio_19 | Precipitation of coldest quarter | mm |
| bdod30_60_cm | Bulk density of fine earth (30–60 cm depth) | kg/dm³ |
| phh2o30_60_cm | Soil pH in H_2_O (30–60 cm depth) | pH |
| nitrogen30_60_cm | Total nitrogen content (30–60 cm depth) | g/kg |
| ocd30_60_cm | Organic carbon density (30–60 cm depth) | kg/m³ |
| soc30_60_cm | Soil organic carbon content (30–60 cm depth) | g/kg |
| cfvo30_60_cm | Coarse fragments volumetric fraction (30–60 cm depth) | cm³/dm³ |
| clay30_60_cm | Clay content (30–60 cm depth) | g/kg |
| sand30_60_cm | Sand content (30–60 cm depth) | g/kg |
| silt30_60_cm | Silt content (30–60 cm depth) | g/kg |
| dem | Elevation | m |
| scd | Soil depth | cm |
| slope | Terrain slope | degrees (°) |
| aspect | Terrain aspect (slope orientation) | degrees (°) |

Collinearity among all environmental predictors was assessed using Pearson correlation coefficients. Variables with pairwise correlation r > 0.9 were considered highly correlated and grouped accordingly.

**Table D2. Groups of highly correlated environmental predictors identified using Pearson correlation (r > 0.9)** Variables retained for modeling are shown in **bold** and were selected based on both the correlation analysis and their ecological relevance for the species.

| *Correlated group 1* | *Correlated group 2* | *Correlated group 3* | *Uncorrelated variables* |
| --- | --- | --- | --- |
| **"bio_1”**  "bio_5"  "bio_6"  "bio_8"  "bio_9"  **"bio_10"**  **"bio_11"** "bdod30_60_cm" "phh2o30_60_cm" | **"bio_12"**  "bio_13"  “bio_14"  "bio_16"  "bio_17"  **"bio_18"**  **"bio_19"** | "nitrogen30_60_cm" "ocd30_60_cm" "soc30_60_cm" "dem"  **"scd"** | "bio_2"  "bio_3"  "bio_4"  "bio_7"  "bio_15" **"cfvo30_60_cm" "clay30_60_cm" "sand30_60_cm" "silt30_60_cm"** **"slope"**  "aspect" |

**Appendix E: European projections of past, current and future abiotic niche suitability continuous maps for *Xatartia scabra***

This appendix provides continuous maps of abiotic niche suitability for *X. scabra* modeled under past, current, and future climatic conditions. Suitability is represented as a gradient from blue (low suitability) to yellow (high suitability). Maps cover the full calibration extent, while the versions displayed in the main article are cropped to the Pyrenees.

Past projections include the Last Glacial Maximum (26,500-17,500 years BP), Heinrich Stadial 1 (17,500-14,700 BP), Bølling-Allerød (14,700-12,900 BP), Younger Dryas (12,900-11,700 BP), Greenlandian (11,700-8,200 BP), Northgrippian (8,200-4,200 BP), and Meghalayan (4,200 BP-present CE), followed by the current baseline period (1950-1981 CE). Future projections are displayed under three Shared Socioeconomic Pathways (SSP126, SSP370, SSP585) for four time intervals: 2010-2040, 2041-2070, and 2071-2100.

**Current period**

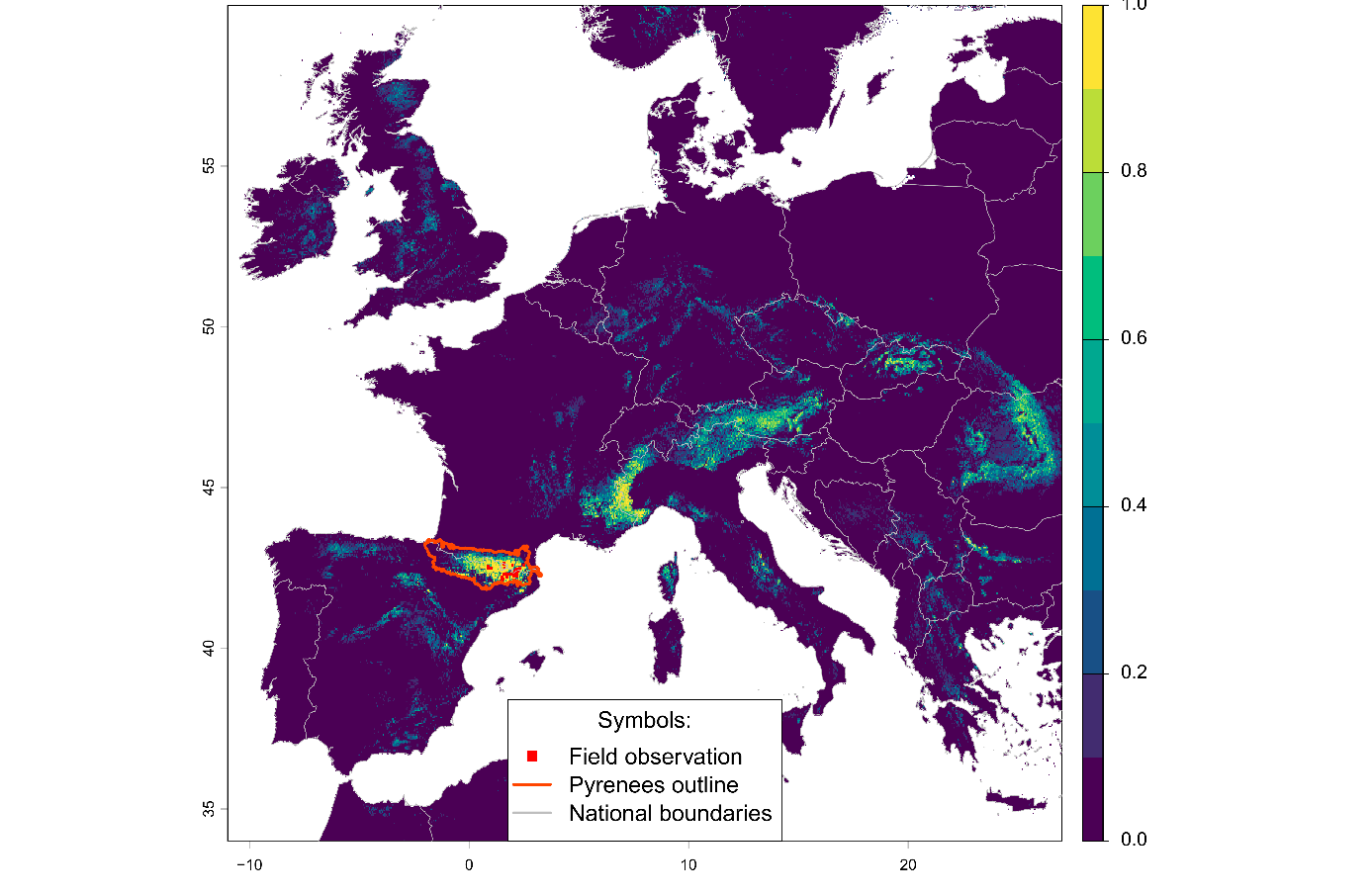

Past projected maps of abiotic suitability for *Xatartia scabra*.

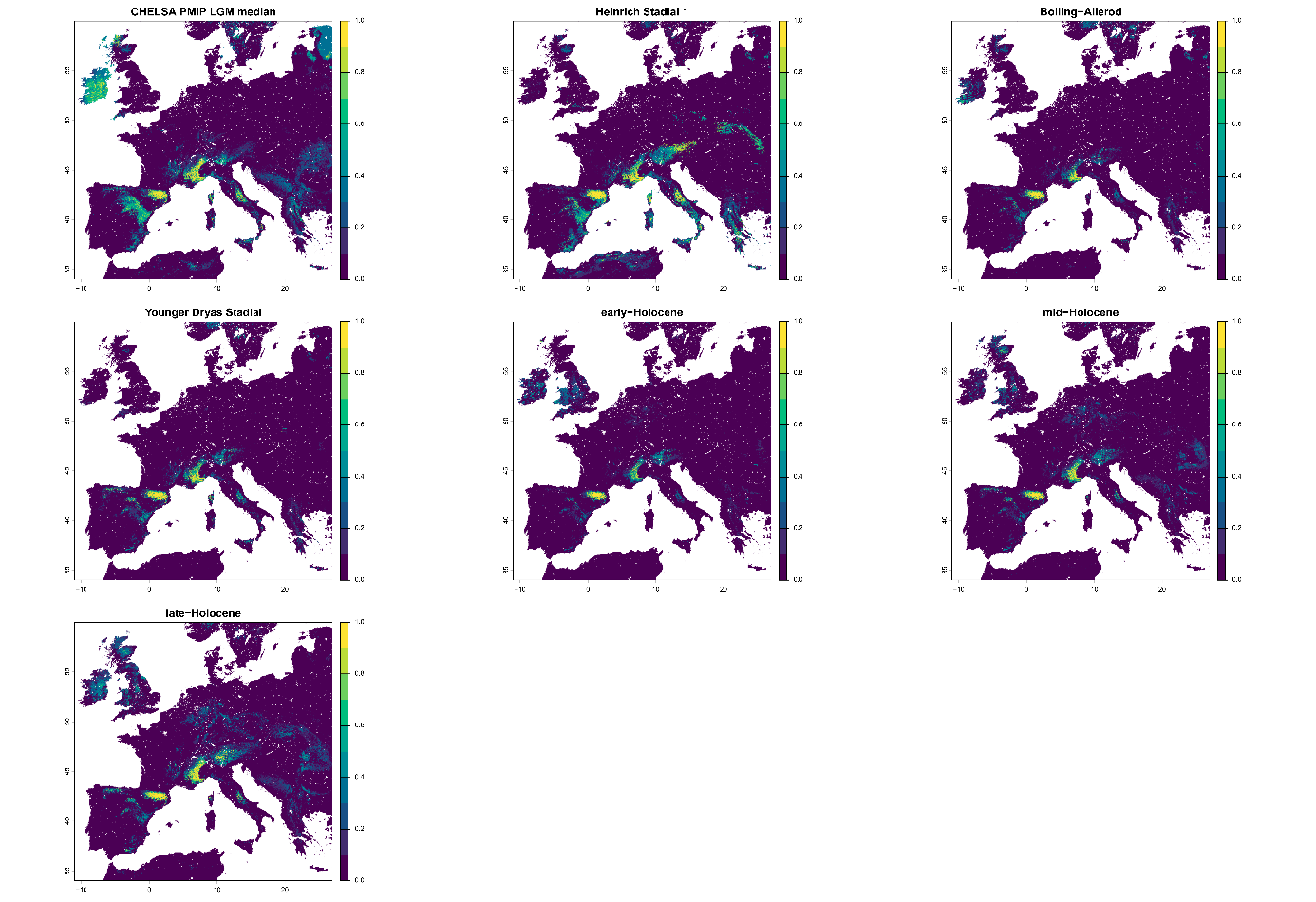

Future projected maps of abiotic suitability for *Xatartia scabra*.

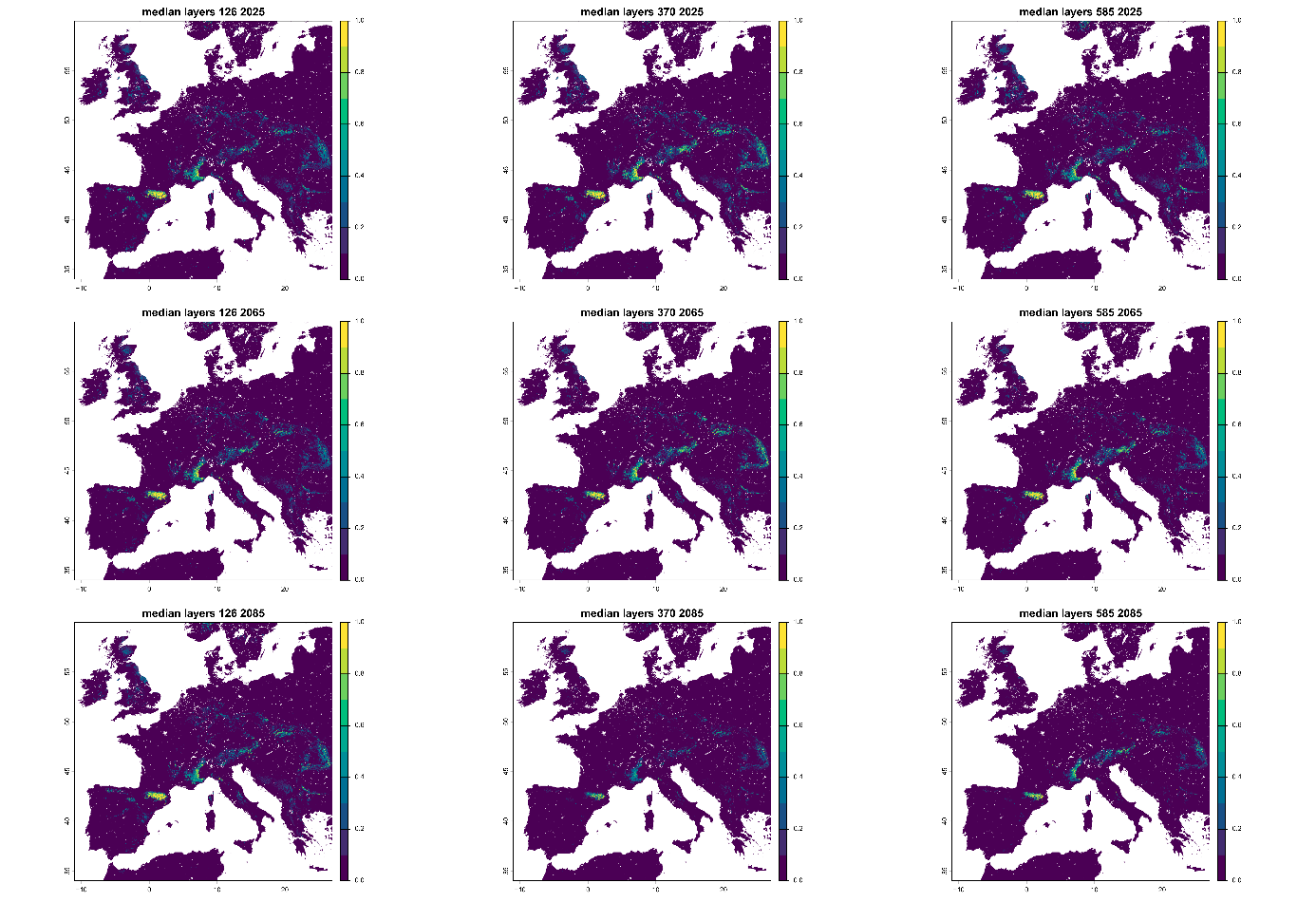

**Appendix F: Missing data rate per individual (%)**

The bar plot displays the percentage of missing data for each individual, colored according to population.

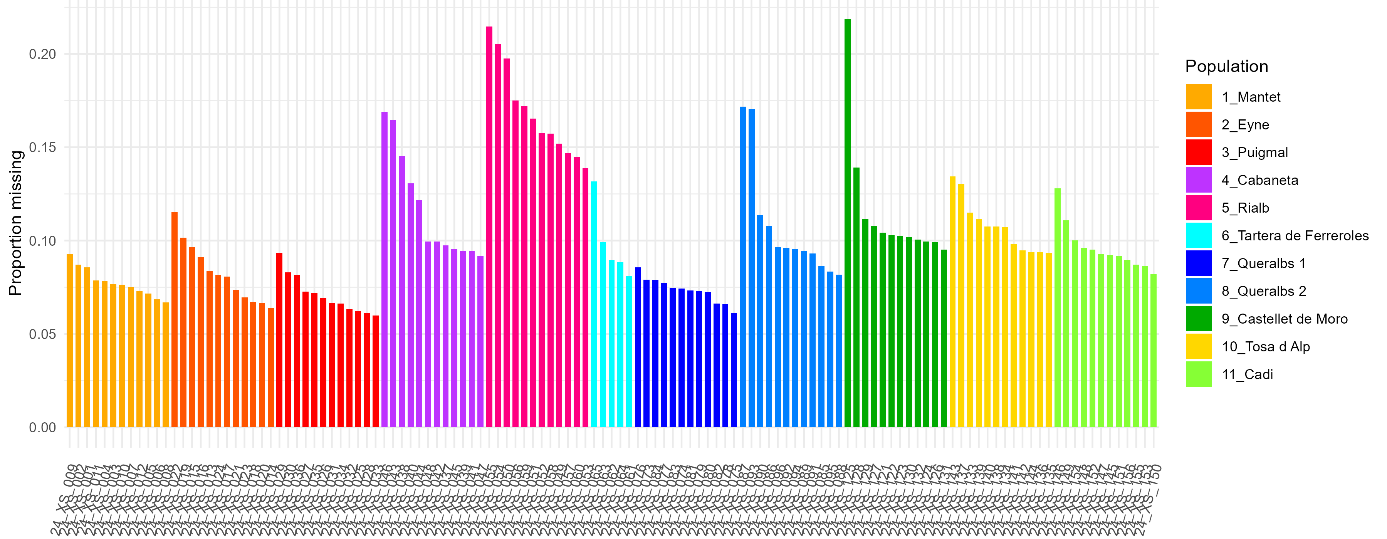

**Appendix G: Cross-entropy criterion as a function of the number of ancestral populations**

The plot shows the cross-entropy values obtained for different numbers of ancestral populations (K = 1–12) using the sNMF algorithm. Cross-entropy decreases sharply from K = 1 to K = 2, indicating a major genetic subdivision in the dataset, and continues to decline until reaching a minimum at K = 6.

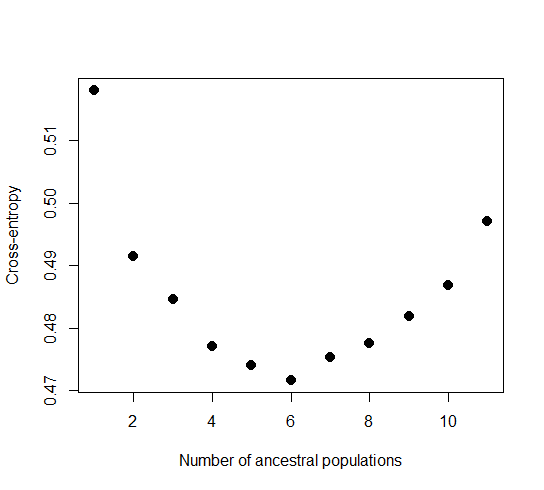

**Appendix H: DILS model fit and parameter estimates**

The parameter settings implemented in the configuration file (config.yaml) to run DILS were as follows: region = noncoding, nspecies = 2, nameA = EAST, nameB = WEST, lightMode = TRUE, useSFS = 1, nameOutgroup = NA, population_growth = constant, modeBarrier = bimodal, max_N_tolerated = 0.1, Lmin = 100, nMin = 6, μ = 7 × 10⁻⁹, ρ/θ = 0.5, N ∈ [500, 100,000], Tsplit ∈ [0, 100,000], and M ∈ [1, 40].

**Table H1: Goodness-of-fit statistics and posterior parameter estimates for three representative DILS runs showing acceptable statistical support and biological consistency.**

|  | **N_min: 500**  **N_max: 75000**  **Tsplit_min: 0**  **Tsplit_max: 150000** | | | | | **N_min: 500**  **N_max: 100000**  **Tsplit_min:0**  **Tsplit_max: 100000** | | | **N_min: 500**  **N_max: 100000**  **Tsplit_min:0**  **Tsplit_max: 50000** | | | |
| --- | --- | --- | --- | --- | --- | --- | --- | --- | --- | --- | --- | --- |
| **stats** | | **mean_exp** | **mean_obs** | **pvals_fdr_corrected** | **mean_exp** | | **mean_obs** | **pvals_fdr_corrected** | | **mean_exp** | **mean_obs** | **pvals_fdr_corrected** |
| sf_avg | | 0 | 0 | **0.5** | 1,00E-05 | | 0 | **0.5** | | 4,00E-05 | 3,00E-05 | **0.38529** |
| sf_std | | 5,00E-05 | 0 | **0.5** | 0.00014 | | 0 | 0 | | 0.00042 | 0.00072 | 0.09425 |
| sxA_avg | | 0.00141 | 0.00143 | **0.49346** | 0.00155 | | 0.00166 | **0.39669** | | 0.00148 | 0.00168 | **0.26477** |
| sxA_std | | 0.00263 | 0.00344 | 0.0039 | 0.00305 | | 0.00375 | **0.1518** | | 0.00297 | 0.00396 | 0.04615 |
| sxB_avg | | 0.00193 | 0.0019 | **0.48783** | 0.00179 | | 0.00184 | **0.42738** | | 0.00141 | 0.0017 | 0.08112 |
| sxB_std | | 0.00315 | 0.00417 | 0 | 0.00329 | | 0.00415 | 0.04006 | | 0.0029 | 0.004 | 0.03744 |
| ss_avg | | 0.00261 | 0.00184 | 0.07718 | 0.0029 | | 0.00182 | 0.04006 | | 0.00213 | 0.0018 | **0.18769** |
| ss_std | | 0.00379 | 0.00421 | **0.13136** | 0.00481 | | 0.00408 | **0.19993** | | 0.00416 | 0.00422 | **0.43723** |
| piA_avg | | 0.00175 | 0.00141 | **0.20924** | 0.00196 | | 0.0015 | **0.18674** | | 0.00159 | 0.00147 | **0.37253** |
| piA_std | | 0.00208 | 0.00248 | 0.00669 | 0.00274 | | 0.00246 | **0.29475** | | 0.00246 | 0.00255 | **0.41524** |
| piB_avg | | 0.00196 | 0.00161 | **0.12591** | 0.00205 | | 0.00161 | **0.16016** | | 0.00156 | 0.00151 | **0.41524** |
| piB_std | | 0.00216 | 0.00268 | 0 | 0.00278 | | 0.00258 | **0.351** | | 0.00245 | 0.00258 | **0.41524** |
| thetaA_avg | | 0.00176 | 0.00143 | **0.21206** | 0.00195 | | 0.00152 | **0.19993** | | 0.00158 | 0.00152 | **0.41524** |
| thetaA_std | | 0.002 | 0.00241 | 0.00603 | 0.00263 | | 0.00247 | **0.35719** | | 0.00238 | 0.00257 | **0.37253** |
| thetaB_avg | | 0.00199 | 0.00164 | **0.12578** | 0.00205 | | 0.0016 | **0.16016** | | 0.00155 | 0.00153 | **0.45925** |
| thetaB_std | | 0.0021 | 0.00264 | 0 | 0.00268 | | 0.00251 | **0.351** | | 0.00236 | 0.00259 | **0.33187** |
| DtajA_avg | | -0.02321 | -0.03199 | **0.4472** | 0.00709 | | -0.02025 | **0.29475** | | 0.00214 | -0.04542 | 0.09425 |
| DtajA_std | | 0.77433 | 0.6855 | 0.05644 | 0.73527 | | 0.70083 | **0.29475** | | 0.68594 | 0.70107 | **0.41524** |
| DtajB_avg | | -0.05231 | -0.03843 | **0.39792** | -0.01005 | | 0.01747 | **0.2756** | | 0.01027 | -0.01937 | **0.21496** |
| DtajB_std | | 0.79635 | 0.72494 | 0.02106 | 0.74638 | | 0.72354 | **0.32509** | | 0.68217 | 0.68817 | **0.46307** |
| divAB_avg | | 0.00192 | 0.00164 | **0.21325** | 0.0021 | | 0.00173 | **0.21002** | | 0.00175 | 0.00167 | **0.41524** |
| divAB_std | | 0.00198 | 0.00263 | 0 | 0.00265 | | 0.00248 | **0.351** | | 0.00245 | 0.00261 | **0.38735** |
| FST_avg | | 0.012 | 0.01415 | **0.32289** | 0.01816 | | 0.02466 | **0.18674** | | 0.03572 | 0.02234 | **0.10764** |
| FST_std | | 0.09042 | 0.07524 | 0.00669 | 0.10137 | | 0.0945 | **0.32457** | | 0.13982 | 0.10538 | 0.08112 |

**Table H2: Posterior parameter estimates and 95% highest posterior density (HPD) intervals for three representative DILS runs with acceptable statistical support and biological consistency** (ABC framework, http://dils.univ-lyon1.fr/ )

Simulation parameter settings: N_min: 500, N_max: 75000, Tsplit_min: 0, Tsplit_max: 150000

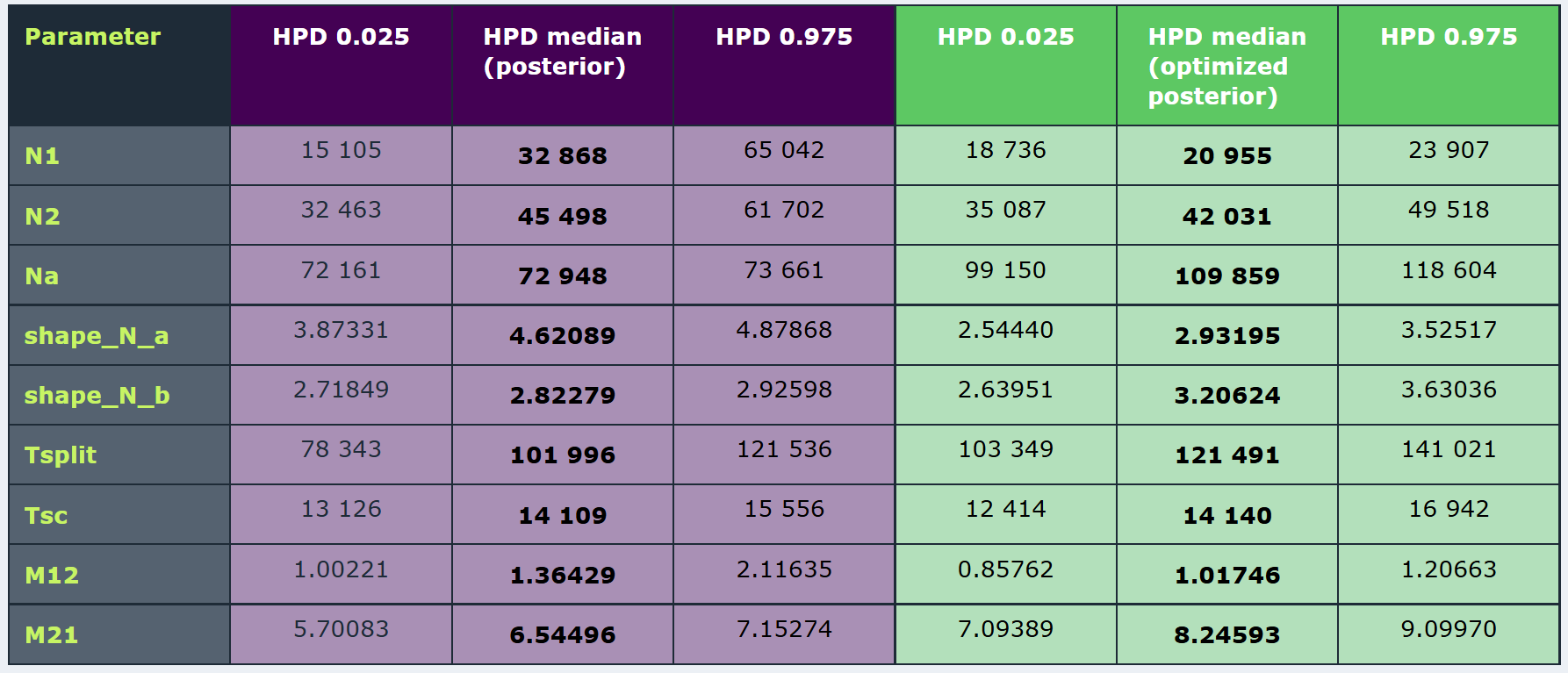

Simulation parameter settings: N_min: 500, N_max: 100000, Tsplit_min:0, Tsplit_max: 100000

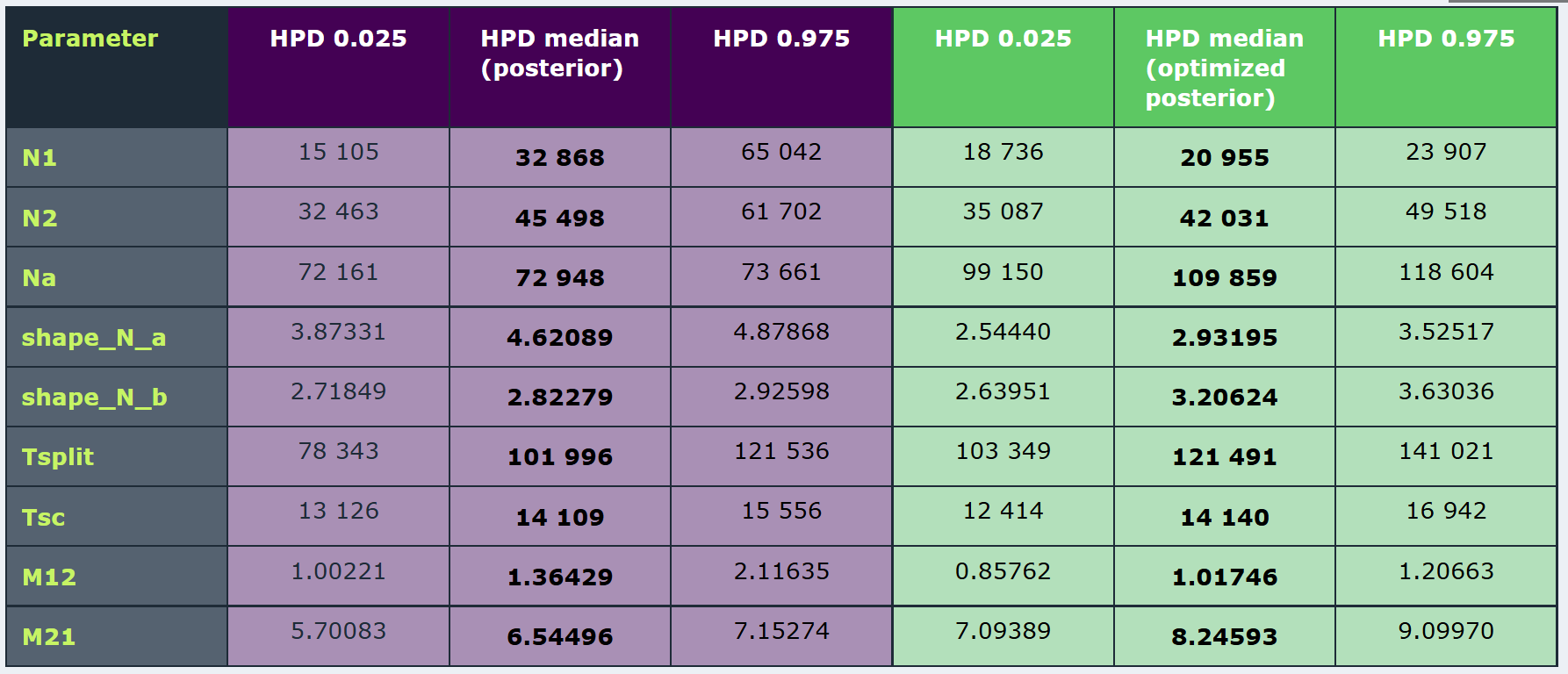

Simulation parameter settings: N_min: 500, N_max: 100000, Tsplit_min:0, Tsplit_max: 50000

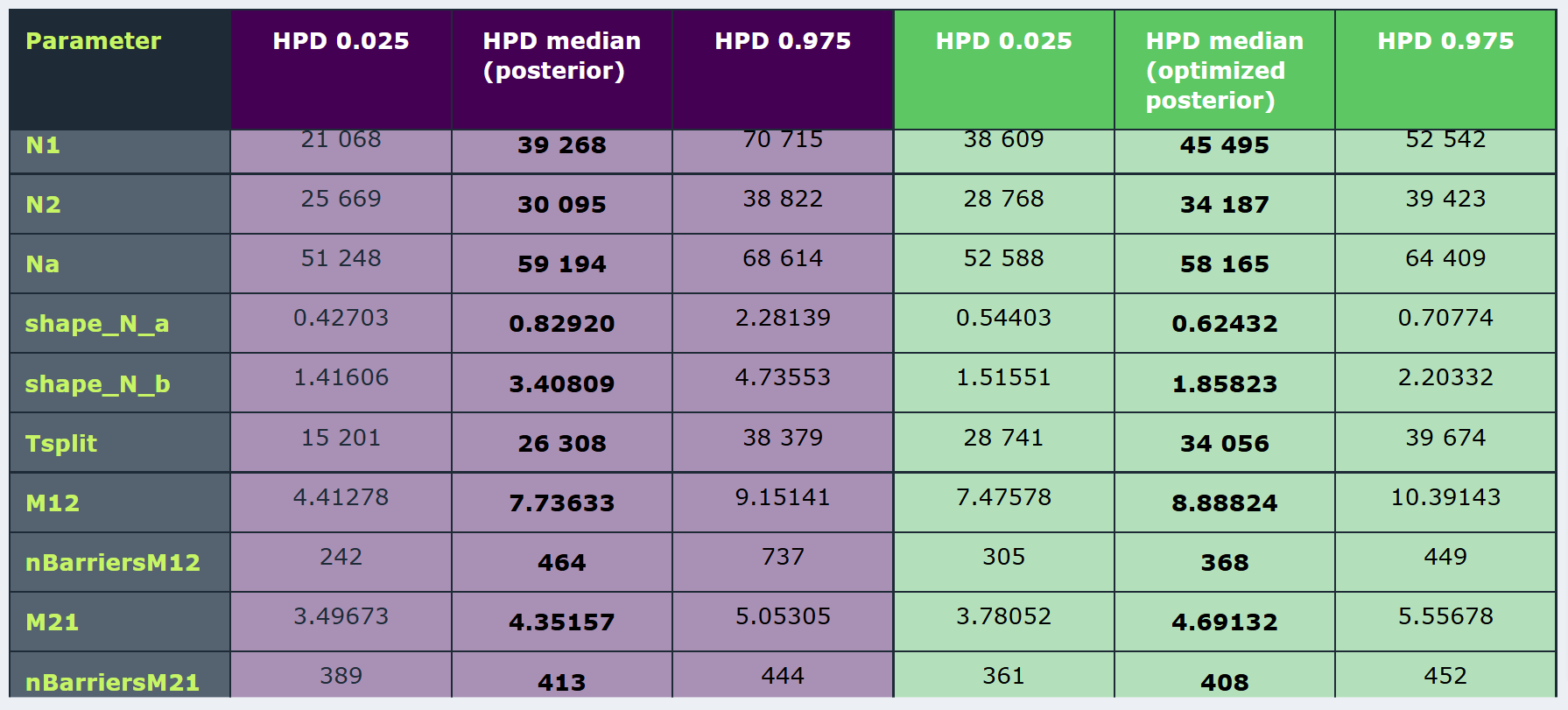
